# Rapid nuclear deadenylation of mammalian messenger RNA

**DOI:** 10.1101/2021.11.16.468655

**Authors:** Jonathan Alles, Ivano Legnini, Maddalena Pacelli, Nikolaus Rajewsky

## Abstract

Poly(A) tails protect RNAs from degradation and their deadenylation rates determine RNA stability. Although poly(A) tails are generated in the nucleus, deadenylation of tails has mostly been investigated within the cytoplasm. Here, we combined long-read sequencing with metabolic labeling, splicing inhibition, and cell fractionation experiments to quantify, separately, the genesis and trimming of nuclear and cytoplasmic tails *in vitro* and *in vivo*. We present evidence for genome-wide, nuclear synthesis of tails longer than 200 nt, which are rapidly shortened within minutes after transcription. Our data show that rapid deadenylation is a nuclear process, and that different classes of transcripts and even transcript isoforms have distinct nuclear tail lengths. For example, many long-noncoding RNAs escape rapid nuclear deadenylation. Modelling deadenylation dynamics predicts nuclear deadenylation about 10 times faster than cytoplasmic deadenylation. In summary, our data suggest that nuclear deadenylation is a key mechanism for regulating mRNA stability, abundance, and subcellular localization.

## Introduction

Poly(A) tails are a hallmark of eukaryotic gene expression and protect RNAs from degradation (Garneau et al., 2007) while promoting translation of mRNAs (Beilharz & Preiss, 2007; Chorghade et al., 2017; Munroe & Jacobsen, 1990; Preiss et al., 1998; Weill et al., 2012), in particular during early development (Palatnik et al., 1984; Sheets et al., 1994; Subtelny et al., 2014). Poly(A) tail length is dynamically regulated (Eckmann et al., 2011) and complete deadenylation typically precedes decapping and exonucleolytic decay.

Poly(A) tails are synthesized post-transcriptionally by Poly(A) Polymerase (PAP) after 3’ end cleavage of the nascent transcript (E. Wahle, 1991). Biochemical studies indicate that PAP is stimulated by the mRNA 3’-end processing factor CPSF (Christofori & Keller, 1989) and the nuclear poly(A) binding protein PABPN1 (Elmar Wahle, 1991), which also acts as a molecular ruler to limit the length of the synthesized tail to ca. 250 nt (Keller et al., 2000; Kühn et al., 2009). These *in vitro* studies are in line with radioactive labeling experiments showing similar poly(A) length profiles of newly synthesized RNA (Brawerman & Diez, 1975; Sheiness et al., 1975). Poly(A) tails are typically shortened throughout an mRNA lifetime and deadenylation rates determine mRNA stability in a gene-specific manner (Brown & Sachs, 1998; Cao & Parker, 2001; Eisen et al., 2020; Sheiness et al., 1975).

mRNA 3’-end processing is tightly coupled to other nuclear processing steps such as splicing and RNA export (Misra & Green, 2016). RNA-sequencing studies identified co-transcriptional splicing as the dominant mode for mammalian RNA processing (Ameur et al., 2011; Tilgner et al., 2012), while a more recent study reports up to 50% of all genes being processed by post-transcriptional splicing, involving complex coordination between individual introns (Coté et al., 2020; Drexler et al., 2020). Splicing of terminal introns is in this context of particular importance, since it is mechanistically linked to pre-mRNA cleavage and polyadenylation (Niwa & Berget, 1991; Vagner et al., 2000) and in some cases used as a mechanism for regulated nuclear retention of transcripts (Bahar Halpern et al., 2015). Splicing of long noncoding RNAs on the other hand is highly complex, generating a multitude of isoforms for individual genes (Deveson et al., 2018). While being processed in the nucleus through the same pathways as messenger RNAs, lncRNAs are in many cases retained in the nucleus (Djebali et al., 2012; Tilgner et al., 2012). Splicing kinetics has been identified as one factor impacting export, but other features such as *cis*-regulatory sequence elements contribute to active retention of lncRNAs in the nucleus (Lubelsky & Ulitsky, 2018; Palazzo & Lee, 2018). Poly(A) tails have been shown to be beneficial but not strictly required for efficient export (Dower et al., 2004; Wilusz et al., 2012).

The CCR4-NOT (Collart, 2016) and PAN2-PAN3 (Wolf & Passmore, 2014) complexes have been identified as major effectors of cytoplasmic mRNA deadenylation in human and yeast. Initial trimming of poly(A) tails has been attributed to PAN2-PAN3 (Brown & Sachs, 1998; Yamashita et al., 2005), while CCR4-NOT was shown to completely deadenylate tails (Yi et al., 2018). Both complexes were found shuttling between cytoplasm and nucleus (Goldstrohm & Wickens, 2008) suggesting that poly(A) turnover may have a nuclear component.

A recent study concludes that mRNAs emerge in the cytoplasm with poly(A) tails of ca. 130-150 nt on average (Eisen et al., 2020), which implies drastic nuclear shortening under the assumption that poly(A) tails are globally synthesized at a length of 250 nt.

In this study, we investigate nuclear processing of poly(A) tails on a genome-wide scale. We first present evidence for genome-wide synthesis of long poly(A) tails by analyzing full-length mRNA and poly(A) sequencing (FLAM-seq) datasets (Legnini et al., 2019). We validate accumulation of long poly(A) tails by splicing inhibition in HeLa cells. Metabolic labeling of newly synthesized RNA in combination with poly(A) profiling indicates rapid global shortening of poly(A) tails within minutes after synthesis. Subcellular fractionation of HeLa cells and mouse brains reveals nuclear deadenylation as a global mechanism for gene specific diversification of poly(A) tail profiles before export. We finally aggregate nuclear and cytoplasmic poly(A) tail length profiles into a quantitative mathematical model of poly(A) tail and expression dynamics which predicts fast nuclear deadenylation of poly(A) tails, with strong regulatory potential.

## Results

### Pre-mRNAs are synthesized with long poly(A) tails

To investigate the poly(A) tail length distribution of nascent transcripts we developed a computational pipeline for extracting unspliced reads from datasets generated by FLAM-seq. We applied stringent criteria to remove ambiguous reads which may not be truly indicative of unspliced transcripts (Methods). We reanalyzed datasets (Legnini et al., 2019) of HeLa cells, induced pluripotent stem cells (iPSCs) and cerebral organoids (Fig. 1A), and detected between 77 and 1313 unspliced (‘intronic’) reads per dataset (Fig. 1B), which corresponded to 0.03 - 0.22% of all reads (Fig. S1A). Notably, iPSCs and organoids had a significantly higher fraction of unspliced reads indicating potential differences in splicing kinetics.

**Fig. 1.**
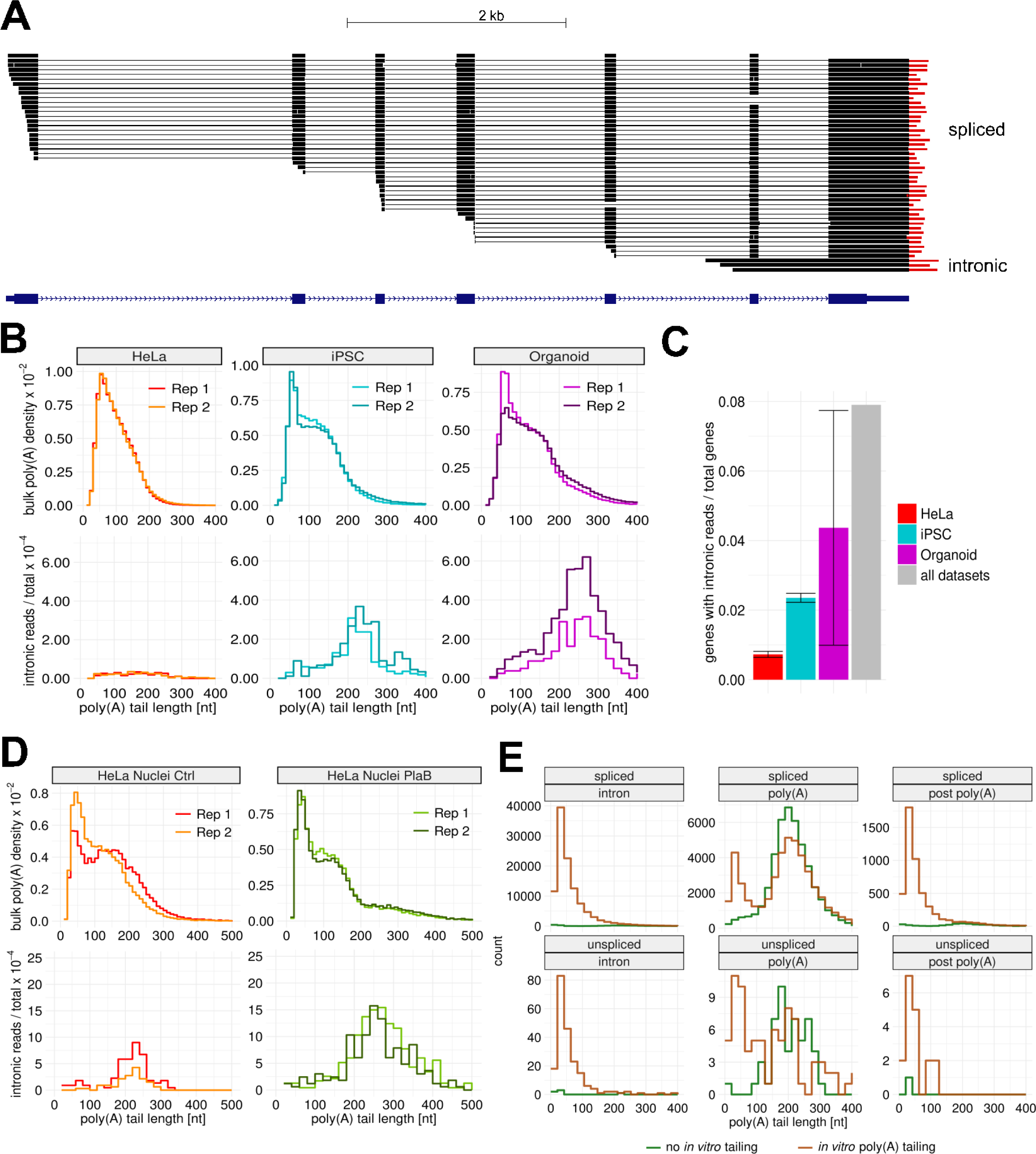
Unspliced RNAs have long poly(A) tails. **A** Browser shot example of detected unspliced read in for gene AHSG in Organoid replicate 1 FLAM-seq dataset (number spliced reads = 317; num unspliced = 3). **B** (TOP) Bulk polyA tail length density distributions for replicates HeLa S3, iPSC, and Organoid datasets in replicates (BOTTOM) Histogram of intronic poly(A) tail length normalized to total reads per dataset for corresponding datasets and replicates. **C** Fraction of genes with detected unspliced, intronic reads normalized to total number of genes detected in each dataset (total number per sample in legend). **D** (TOP) bulk poly(A) tail length density distribution of nuclei from HeLa S3 nuclei control and PlaB experiments for replicate datasets. (BOTTOM) histogram of intronic poly(A) tail length normalized to total read counts per dataset for HeLa S3 nuclei control and PlaB splicing inhibition. Number of reads per dataset/replicate for bulk/intronic are displayed inside legend. **E** Poly(A) tail length distributions for Nanopore direct RNA sequencing (Drexler et al.) reads grouped by annotated as intronic, poly(A) site, or post poly(A) site reads for sequencing protocols involving enzymatic poly(A) tailing of RNA or no tailing. Reads were categorized as spliced (TOP) or unspliced (BOTTOM) by our computational pipeline.

Since differences in read length could be linked to different errors in poly(A) quantification, we recorded the raw read length distributions for bulk and intronic reads (Fig. S1B). These distributions were very similar across replicate experiments, suggesting consistent poly(A) tail length estimates between spliced and unspliced reads.

Intronic poly(A) tail length distributions were longer than bulk poly(A) distributions in each dataset with a peak at 208 nt for iPSCs and 232 nt for organoids. HeLa datasets also showed longer poly(A) tails for intronic reads (median 151 nt) but with a higher contribution of poly(A) tails shorter than 150 nt, which might be an artefact of transcripts with retained introns, rather than *bona fide* nascent RNAs.

To investigate whether the detected unspliced, polyadenylated transcripts were representative of transcriptome-wide nascent RNA processing patterns, we first estimated the fraction of genes for which unspliced reads were detected. We found that between 1% and 7% of all genes per dataset had detectable unspliced reads and this number increased to 8% when merging all intronic reads against a background of all detected genes in each dataset (Fig. 1C). We found little overlap in detected genes with intronic reads between datasets (Fig. S1C), which hinted at an unbiased, random sampling process.

Since our pipeline excluded genes if their introns overlapped with coding or 3’UTR sequences of other isoforms, we quantified the impact of this restriction on our analysis: 49-53% of total detected genes in each dataset had unambiguous intron annotations (Fig. S1D), which defined the upper bound of detectable genes with unspliced reads taking into account the average length of FLAM-seq reads.

We downsampled intronic reads and each computed the fraction of genes with intronic reads against all detected genes in a dataset. This analysis revealed a nearly linear relationship, showing that higher sequencing depth would further increase the number of genes with intronic reads (Fig. S1E). We further excluded additional potential biases in our analysis by comparing other features of intronic reads such as intron length (Fig. S1F-G) and gene expression levels (Fig. S1H-I).

In summary, we developed a computational pipeline which identifies poly(A) tail length of unspliced transcripts. Our data indicate that unspliced polyadenylated RNAs have poly(A) tails of more than 200 nt in length on average.

### Splicing inhibition causes accumulation of unspliced RNA with long tails

To validate our analysis of unspliced RNAs in FLAM-seq datasets, we treated HeLa cells for 3 hours with the SF3b inhibitor Pladeinolide B (PlaB), which blocks spliceosome assembly on pre-mRNA (Martínez-Montiel et al., 2016). To enrich unspliced RNAs, we isolated nuclei from control and PlaB-treated cells. We observed a more than tenfold increase in unspliced reads in RNA extracted from nuclei compared to RNA extracted from whole cells, validating the enrichment of pre-mRNA in nuclear preparations. We found a 3-fold increase in intronic reads upon splicing inhibition (from 0.4% to 1.3% of total reads, Fig. S2A). We detected 54-304 intronic reads per replicate (Fig. 1D) and for 1% of total genes in control and 3.5% in PlaB-treated cells (Fig. S2B). We again found little overlap of genes with intronic reads between individual replicates, arguing against biases in detecting unspliced transcripts (Fig. S2C).

Poly(A) tail length distributions of intronic reads had a median of 205 nt in control and 242 nt in PlaB-treated HeLa nuclei, with a marked reduction of unspliced RNAs with tails shorter than 150 nt compared to bulk RNA preparations from HeLa cells (Fig. 1B). We binned genes by expression and investigated the poly(A) tail length of intronic reads for each bin. We did not observe expression-dependent differences in intronic poly(A) length but noticed a significant increase in poly(A) length between intronic reads from control and PlaB conditions (Fig. S2D). Consistent with results from bulk RNA preparations, the relative fractions of unspliced reads decreased with higher gene expression for control samples. This trend was less prominent for PlaB-treated HeLa cells which may indicate differences in the efficacy of PlaB in blocking splicing for different classes of genes (Fig. S2E).

Comparing the bulk poly(A) length distributions revealed global shortening of poly(A) tail length of about 11 nt upon PlaB treatment (Fig. S2F). At the same time, we noticed an accumulation of poly(A) tails longer than 200 nt upon PlaB treatment which may be explained by accumulation of unspliced reads which were too short to cover terminal introns (‘false negative’ unspliced reads). Differences in poly(A) length and fold-changes in expression between control and PlaB were not correlated (Fig. S2G), but we found several genes for which changes in gene expression coincided with poly(A) tail extension (MYC, IER3, KLF5, RBMX) or with tail shortening (MAT2A, STX10).Transcripts with increasing poly(A) tail length upon PlaB treatment were typically less stable (Fig. S2H) and had longer 3’UTRs (Fig. S2I).

We further compared our analysis to a recently published Nanopore direct RNA sequencing datasets of nascent, chromatin-associated RNA (Drexler et al., 2020). In this study, non-polyadenylated RNA is analyzed by *in vitro* A-tailing of RNA using an *E.Coli* polyA polymerase before library preparation and subsequent Nanopore long-read direct RNA sequencing (Workman et al. 2019). Nanopore reads were (separately for spliced and unspliced reads) sorted into three groups, (1) reads mapping upstream of the poly(A) site (2) reads ending at the poly(A) site and (3) reads that extended downstream of the poly(A) site (Fig. 1E). The experimental approach without poly(A) tailing enriched polyadenylated transcripts with long poly(A) tails both for spliced and unspliced reads which ended at annotated polyadenylation sites. As expected, unspliced reads had long poly(A) tails of ca. 200 nt which also validated the poly(A) profiles of intronic reads identified in our FLAM-seq datasets. Poly(A) tailing enriched for reads ending in introns which had poly(A) tails with a median length of 20-35 nt, which were likely the product of the *in vitro* tailing by *E. coli* poly(A) polymerase used to generate these data. In these samples, poly(A) tails are short for all reads except those ending in a poly(A) site, which indicated absence of polyadenylation for nascent transcripts which are not cleaved. Both spliced and unspliced reads, which ended at a poly(A) site, had bimodal poly(A) length distributions which could be interpreted as (pre-)mRNAs having either long or no poly(A) tails (Fig. 1E).

In summary, we validated that our pipeline captured unspliced pre-mRNAs by inhibiting splicing using PlaB and observed global shortening of poly(A) tails but elongated tails of unspliced transcripts. We further analyzed previously published data and showed that poly(A) profiles of unspliced reads were bimodal in non-poly(A) selected chromatin fractions indicating that transcripts were either not polyadenylated or had ca. 200 nt-long tails at the poly(A) site.

### Metabolic labeling reveals poly(A) tail length profiles of newly transcribed RNA

Since poly(A) tail length profiles of unspliced reads were significantly longer than steady-state distributions we investigated the deadenylation kinetics on a genome-wide scale using metabolic labeling of RNA in combination with FLAM-seq (Fig. 2A). We first performed 4-thiouridine (4sU) labeling of HEK cells followed by biotinylation of 4sU moieties and separation of labeled pulldown and supernatant fractions using streptavidin-coated beads (Dölken et al., 2008; Rabani et al., 2011).

**Fig. 2.**
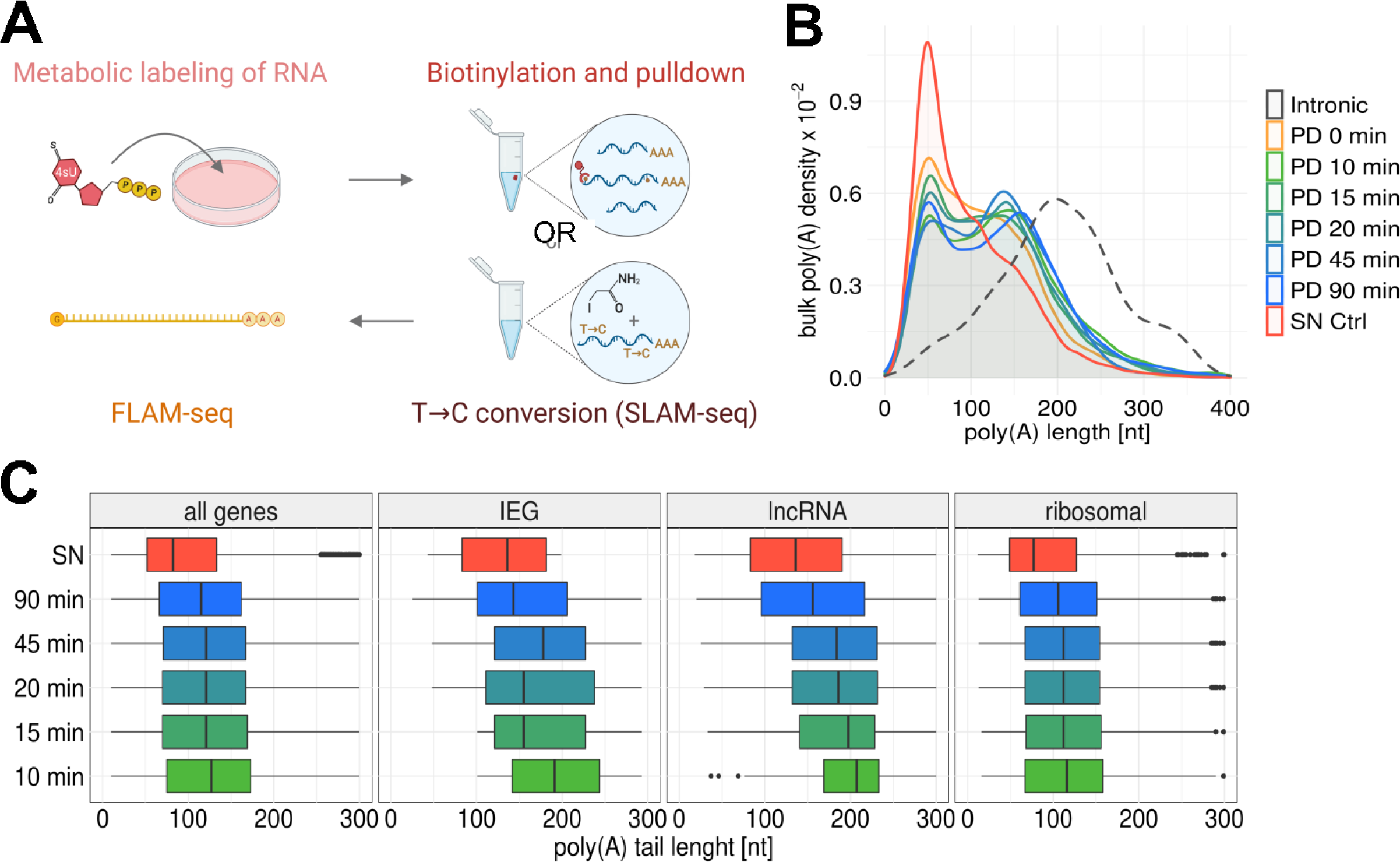
Metabolic labeling indicates rapid shortening of poly(A) tails after synthesis. **A** Experimental outline for 4sU pulldown and SLAM-Seq metabolic labeling experiments. **B** Poly(A) tail length distributions for supernatant (SN) and pulldown (PD) fractions after metabolic labeling with 4sU for indicated time points. Dashed distribution indicates poly(A) tail length for intronic reads from HEK metabolic labeling experiments. **C** Poly(A) tail length distributions of all genes, immediate early genes (IEGs), long noncoding RNAs (lncRNA) and ribosomal protein encoding genes (ribosomal) for different labeling timepoints in 4-SU pulldown experiments.

We labeled HEK cells for up to 90 minutes using 4sU at high concentrations (1 mM) to obtain sufficient labeled RNA for performing FLAM-seq. RNA concentrations obtained after pulldown were proportional to labeling time (Fig. S3A). Comparing pulldown and supernatant concentrations, 1% - 10% of the total RNA pool was captured in pulldown fractions, depending on labeling period, which was in agreement with previous studies (Dölken et al., 2008). We noticed a small fraction of RNA binding to streptavidin beads in control samples without 4sU labeling (0 min), which was also reported before (Dölken et al., 2008). Dot Blots of total RNA before pulldown showed efficient biotinylation and signal intensities proportional to labeling time (Fig. S3B).

In the FLAM-seq data, we observed longer poly(A) tails in pulldown fractions compared to supernatant (Fig. 2B), with a bimodal distribution peaking at 50 nt and 150 nt. The supernatant distribution had only a single mode at 50 nt. Differences in poly(A) tail length profiles between individual labeling time points were minor, indicating little global poly(A) length dynamics throughout the investigated time course.

We noticed an increased poly(A) tail length in the library prepared from a 0 min control experiment, which could indicate preferential binding of longer tails to streptavidin beads in the absence of labeled RNA. Supernatant samples for individual time points were reproducible and indicated absence of systematic biases in poly(A) tail length profiling between time points (Fig. S3C). Comparing transcripts of mitochondrial genes, we observed median lengths of 50 nt in pulldown and supernatant fractions which ensured accurate quantification (Temperley et al., 2010). As mitochondrial poly(A) tail length profiles were similar between pulldown and supernatant fractions independent of labeling period, we concluded that progressive deadenylation is not occurring for mitochondrial transcripts, which had been suggested before (Temperley et al., 2003). Pulldown samples contained the majority of unspliced reads (Fig. S3E), which was expected given that splicing is typically completed within minutes (Rabani et al., 2014).

Poly(A) tail length of unspliced reads peaked at 200 nt, which validated accurate quantification of long poly(A) tails in pulldown experiments (Fig. S3F). We computed differences in median poly(A) tail length per gene between pulldown and supernatant fractions (Fig. S3G) finding that poly(A) tails were 24-32 nt longer in pulldown fractions. Comparing poly(A) tail length profiles for different classes of genes, such as genes encoding ribosomal proteins, immediate early genes (IEGs), or lncRNAs, we found stark differences in their poly(A) tail profiles over the labeling time course (Fig. 2C). lncRNAs and IEGs had long poly(A) tails of around 200 nt after 10 min labeling, whereas tails for ribosomal proteins were on average 120 nt. Differences in poly(A) profiles for individual time points were less pronounced for ribosomal transcripts than for IEGs.

As an orthologous approach for profiling poly(A) tail lengths kinetics we applied SLAM-Seq (Herzog et al., 2017) in combination with long read sequencing and poly(A) quantification. We were able to successfully discern labeled and unlabeled reads and quantified poly(A) tail length of newly synthesized transcripts after 90 and 180 minutes of labeling (Fig. S3H). Comparing labeled and unlabeled reads we observed that newly synthesized RNAs had longer poly(A) tails than pre-existing transcripts, which was consistent with the pulldown experiments. We noticed several technical challenges which affected the comparison of poly(A) profiles between time points and refer to the Supplementary Discussion for an in-depth assessment.

In summary, metabolic labeling experiments indicated that, even for short labeling periods, poly(A) tails of newly synthesized RNAs were significantly shorter than 200 nt for most transcripts. Thus, assuming that mRNAs are synthesized with long poly(A) tails, our data suggest rapid poly(A) deadenylation within minutes after the completion of transcription.

### Poly(A) tails are deadenylated in the nucleus

Our metabolic labeling experiments indicated rapid shortening of poly(A) tails within minutes after synthesis. This raised the question whether initial deadenylation occurs already in the nucleus or after mRNA has been exported, which has been reported to operate on comparable timescales (Chen & van Steensel, 2017; Mor et al., 2010).

We addressed this question by investigating poly(A) tail length profiles in subcellular compartments *in vivo* and *in vitro*. We first performed biochemical fractionation of HeLa cells into cytoplasmic, nucleoplasmic and chromatin fractions (Fig. 3A). To account for possible experimental variability, we produced 6 biological times 2 technical replicates. We evaluated the purity and absence of contamination for each sample by Western Blot (Fig. S4A) and by quantifying the fraction of mitochondrial transcripts (Fig. S4B) and intronic reads (Fig. S4C). We did not find evidence for cytoplasmic contamination, although the fraction of mitochondrial and intronic reads differed between biological replicates. Comparing median poly(A) tail length (Fig. S4D) and gene expression counts (Fig. S4E) further showed high reproducibility between technical replicates.

**Fig. 3.**
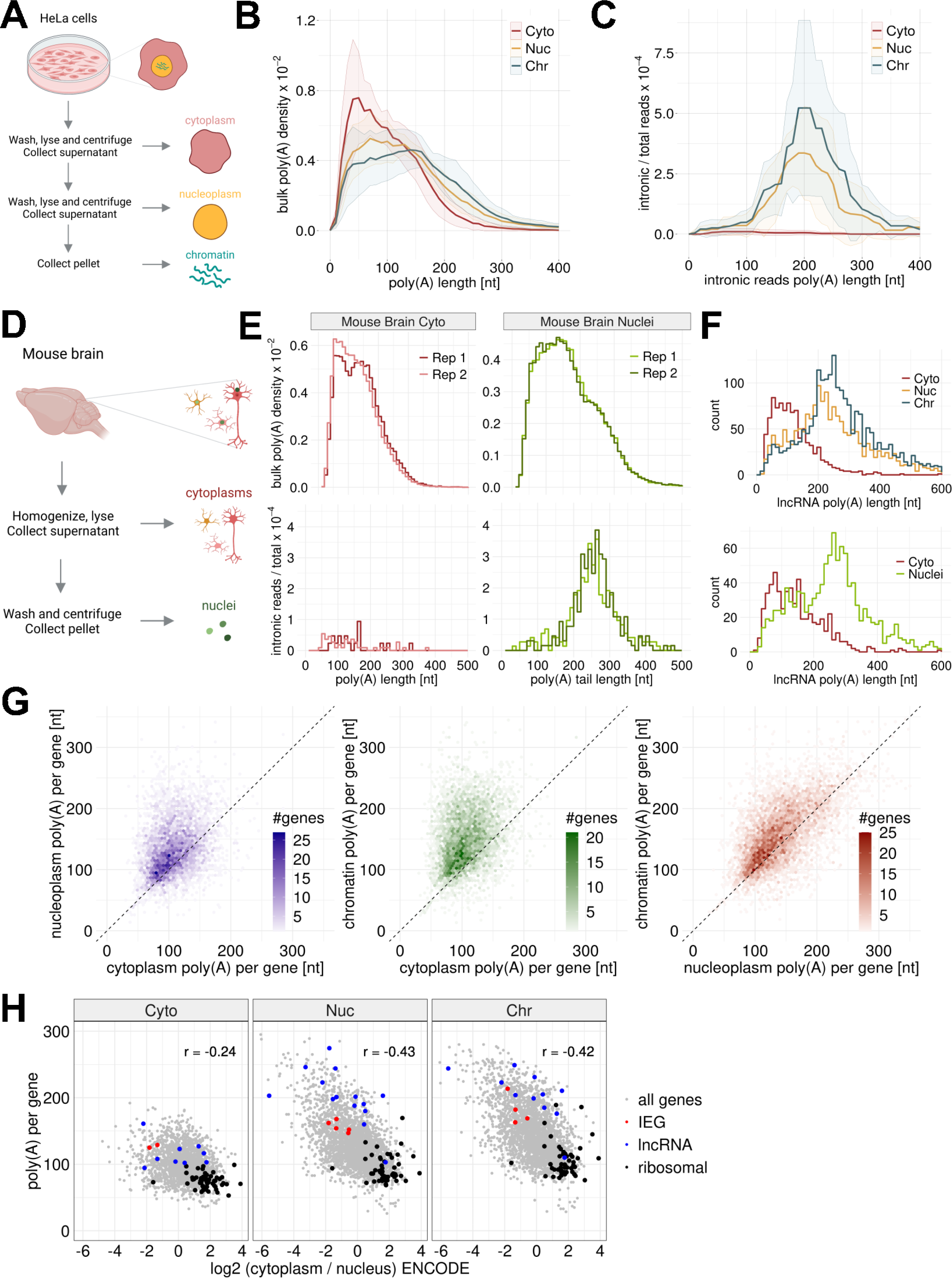
Biochemical fractionation reveals nuclear deadenylation. **A** Experimental outline for biochemical fractionation experiments from HeLa S3 cell lines in cytoplasmic, nucleoplasm and chromatin fractions. FLAM-seq libraries were generated from RNA extracted from fractions. **B** Poly(A) tail length density distributions of cytoplasmic (Cyto), nucleoplasmic (Nuc) and chromatin (Chr) fractions from HeLa S3 cells. Ribbons indicate standard deviation of 12 replicate samples. **C** Poly(A) tail length profiles of intronic reads in HeLa S3 fractions. Ribbons indicate standard deviation of 12 replicate samples. **D** Experimental outline for fractionation of mouse brains into nuclei and cytoplasmic fractions using a Dounce homogenizer. FLAM-seq libraries were generated from RNA extracted from fractions. **E** Poly(A) tail length profiles of mouse brain cytoplasm and nuclei fractions. Bulk distributions include all reads, intronic distributions include unspliced reads. **F** Poly(A) tail length distributions for lncRNAs in HeLa S3 (top) and mouse brain (bottom) fractionation experiments. **G** Comparison of median poly(A) tail length per gene profiles between cellular fractions. **H** Comparison of gene enrichment between nucleus and cytoplasm (inferred from HeLa ENCODE data) and median poly(A) tail length per gene in each fraction.

Comparing length profiles for subcellular fractions indicated variability in poly(A) tail length profiles between biological replicates, while technical replicates were in good agreement (Fig. S4G). To define experimental parameters which could explain the observed differences between biological replicates we fitted a linear model using variables such as the fraction of intronic reads, RNA concentration and others (Methods) to predict the median poly(A) length for each sample. Median poly(A) length was best explained by a scaling factor for each biological replicate, which described the variation from the average poly(A) tail profile over all replicates. Scaling poly(A) tail length profiles accordingly reduces the observed between biological replicates (Fig. S4H). Global biases poly(A) tail length quantification using FLAM-seq could be caused by differences in PCR cycles, input RNA amounts or efficiency of poly(A) selection. We further assumed that the large number of independent replicates for HeLa fractionation experiments accurately accounts for the variability known to be inherent to biochemical fractionation techniques (Jadot et al., 2017). Bulk poly(A) tail length distributions of HeLa chromatin, nucleoplasm and cytoplasm showed progressively shorter poly(A) tails with a median length of 134 nt, 117 nt and 80 nt respectively (Fig. 3B). Poly(A) tail length profiles of intronic reads in chromatin and nucleoplasmic fractions had a median length of 205 nt (Fig. 3C), which was in line with poly(A) tail profiles of unspliced reads in whole cell extracts (Fig. 1B-E). Intronic reads in cytoplasmic fractions were rare (0.02% of total reads vs. 0.6% for nuclear fractions) and had shorter tails of 130 nt. Comparing intronic to bulk profiles hence suggested a rapid shortening process already in the nuclear fractions. We did not find substantial differences in poly(A) tail profiles of genes with detected unspliced transcripts versus those without (Fig. S5A), illustrating that nuclear shortening is a global mechanism.

To investigate molecular properties differentiating genes in their deadenylation dynamics, we compared gene expression by poly(A) tail length bins between fractions (Fig. S5B). As shown before (Chang et al., 2014; Legnini et al., 2019; Lima et al., 2017; Subtelny et al., 2014), highly expressed genes tended to have shorter poly(A) tails in cytoplasmic fractions, a trend which was also seen in nucleoplasm and chromatin but shifted towards longer poly(A) tails. To ensure that nuclear deadenylation is starting from a uniform initial length distribution at the points of synthesis, we compared poly(A) tail length of intronic reads for genes binned by median bulk poly(A) tail length in chromatin and nucleoplasm and observed comparable intronic poly(A) length of ca. 200 nt for each bin (Fig. S5C). We found that short-lived RNAs had longer tails in cytoplasm and a similar trend was seen in nucleoplasm and chromatin (Fig. S5D). 3’UTR length increased proportionally to poly(A) tail length both in cytoplasmic and nuclear fractions (Fig. S5E). We noticed slightly shorter 3’UTRs for nuclear compared to cytoplasmic transcripts, except for those with very short poly(A) tails of less than 50 nt.

We grouped 3’UTR isoforms by shorter proximal and longer distal isoforms and observed isoform-specific deadenylation (Fig. S5F). In summary, the molecular features differentiating poly(A) tail length in the cytoplasm are also associated with poly(A) tail length in nuclear fractions.

We validated nuclear deadenylation *in vivo* by performing fractionation of a mouse brain into cytoplasm and nuclei (Fig. 3D). We showed absence of cytoplasmic contamination in mouse brain nuclei by western blot and analysis of mitochondrial and intronic read fractions in each fraction (Fig. S4A-C). When comparing poly(A) tail length distributions of all reads, we found that nuclear tails were longer than cytoplasmic tails (119 nt vs. 146 nt) but markedly shorter than unspliced intronic reads in the nuclear fraction (Fig. 3E). Unspliced reads in cytoplasmic fractions had shorter poly(A) tails and could reflect transcripts with retained introns as observed for the HeLa cytoplasm. Inspecting poly(A) tail length of different gene biotypes in HeLa and mouse brains, we found that lncRNAs had overall much longer poly(A) tails in the nuclear fractions, both in HeLa and mouse brain samples (Fig. 3F). We found 50-60% of all nuclear lncRNAs with a median poly(A) tail length longer than 200 nt which indicated absence of nuclear deadenylation for those transcripts.

We next compared median poly(A) tail length per gene between fractions (Fig. 3G). Chromatin and nuclear poly(A) tails were on average longer than cytoplasmic tails and we observed a larger dynamic range in nuclear compartments. Chromatin tails were longer than nucleoplasmic tails with a minor offset between fractions. A similar trend was observed when comparing mouse brain cytoplasm and nuclei (Fig. S5H). Different classes of genes showed distinct poly(A) tail length profiles, as for instance observed for ribosomal protein genes which had short poly(A) tails in the nucleus, whereas tails were markedly longer for immediate early genes (IEGs) and lncRNA poly(A) tails (Fig. S5I).

We finally investigated poly(A) tail length in the context of transcript distributions between nucleus and cytosol. We obtained cytoplasmic-to-nuclear transcript ratios from ENCODE RNA-seq data for HeLa cell fractions (Yi et al., 2018) which we found reflective of the ratios inferred from FLAM-seq fractionation data (Fig. S5J). 30% of genes were enriched more than 1.5-fold in the nucleus, which was in line with previous studies investigating subcellular transcript distributions (Halpern et al., 2015). We compared cytoplasmic-to-nuclear enrichments with median poly(A) tail length per gene for each HeLa fraction and found that nuclear tail length is more predictive of transcript enrichment than cytoplasmic tail length (Fig. 3H). Transcripts enriched in the cytoplasm tended to have short tails in all fractions, as illustrated for ribosomal protein genes. On the other hand, lncRNAs highly enriched in the nucleus had long poly(A) tails, while IEGs had intermediate tail length.

In summary, cellular fractionation experiments revealed nuclear deadenylation of poly(A) tails as a gene-specific process *in vitro* and *in vivo*.

### Mathematical modeling of nuclear and cytoplasmic poly(A) tail dynamics predicts fast initial deadenylation

We conceived a simple mathematical model describing mRNA abundance and tail length in the nucleus and the cytosol of a cell to quantitatively address the hypothesis that poly(A) tails are rapidly deadenylated in the nucleus. We described the steady state expression levels of mRNAs in the two compartments as a function of gains and losses, defined by transcription, export, nuclear/cytoplasmic deadenylation and decay. Each process was defined by a rate, which was initially estimated by building a normal distribution around an informed guess (from published data, Fig. S6A, Table S1 and Computational Methods). Rates were then inferred by minimizing the square root differences of simulated mRNA abundances and cytoplasmic and nuclear poly(A) tail length profiles measured by FLAM-seq (Fig. 4A). We observed that the simplest model, which does not allow for nuclear decay and defines a constant export rate, resulted in a buildup of completely deadenylated mRNAs and a bulk distribution of long tails in the nucleus, which did not fit our data (Fig. 4B). Instead, allowing for nuclear decay, or alternatively, setting an export rate dependent on poly(A) tail length, resulted in a better fit for both nuclear and cytoplasmic mRNA distributions (Fig. 4B). We varied some of the model assumptions, including potential contamination of nuclear fractions in fractionation experiments, allowing for nuclear decay or tail-dependent export (Fig. S6B and S6C), and independently estimated the model parameters from individual replicates (Fig. S7A). We observed that our best estimates for the nuclear deadenylation rates were consistently higher than cytoplasmic deadenylation rates (approximately one order of magnitude - ca. 4 nt/min versus 0.5 nt/min, Fig. 4C and Table S1). Nuclear deadenylation may represent an additional, robust layer for global mRNA regulation: we predicted that varying the deadenylation rate in the nucleus results in strong changes in cytosolic mRNA levels (Fig. S7B). In summary, our quantitative modeling approach predicted nuclear deadenylation kinetics which are an order of magnitude faster than in the cytoplasm, along with additional features of nuclear processing such as nuclear decay and tail-length dependent export rates (Fig. 4D).

**Fig. 4.**
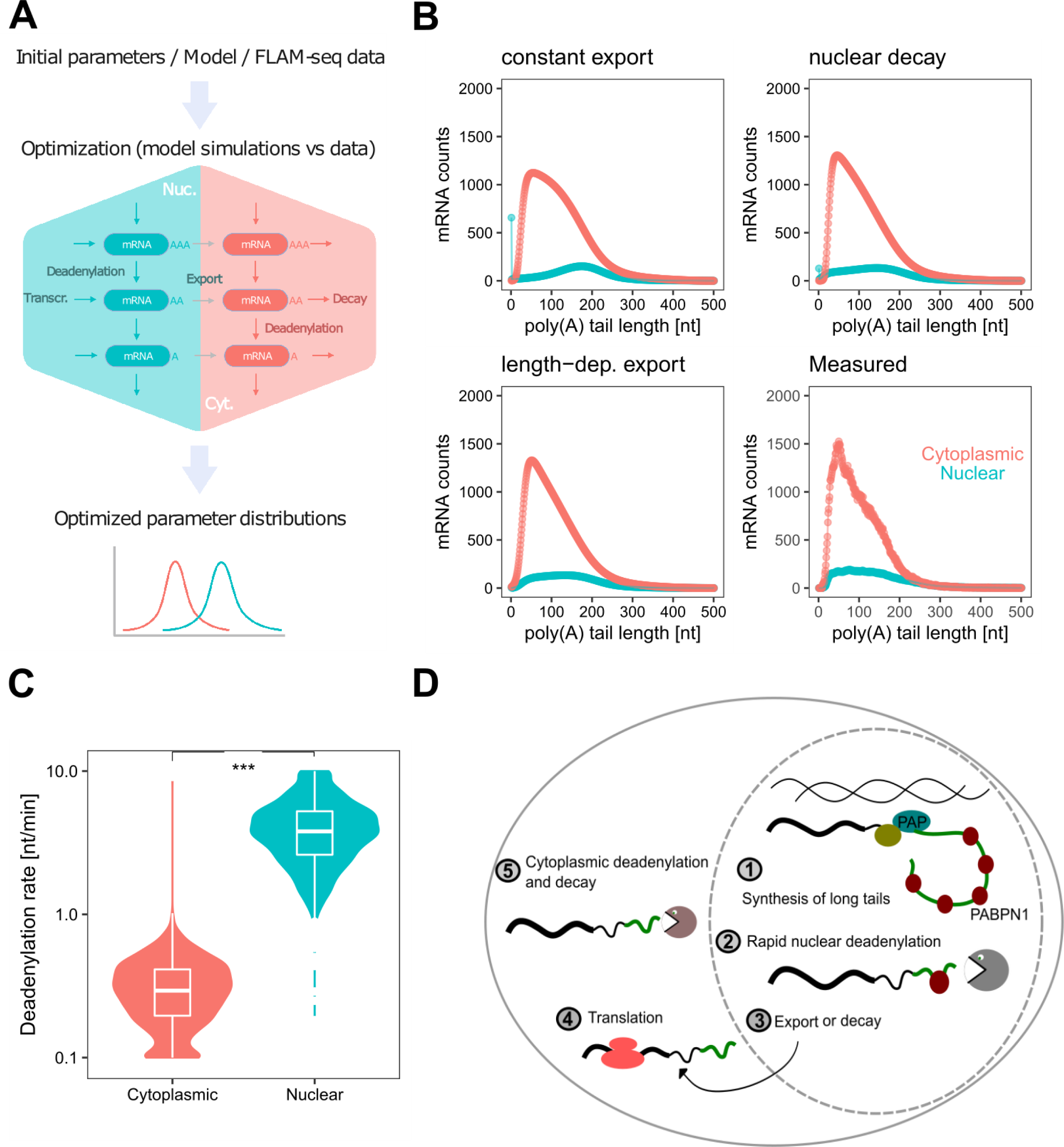
A mathematical model predicts fast nuclear deadenylation. **A** Model description (see methods). **B** Simulations of mRNA levels in the nucleus (green) and cytosol (red) with the basic model assuming constant export (top left), the model assuming constant export and nuclear decay (ndce model; top right), the model assuming tail-dependent export (nndle model; bottom left), compared to the merged FLAM-seq data from 12 fractionation replicates (bottom right). The average of the estimated parameter distributions was used as input. Cytoplasmic contamination in the nuclear fraction of 0% was assumed. **C** Distributions of cytoplasmic (red) and nuclear (green) deadenylation rates, estimated with models assuming nuclear decay (ndce) or tail-dependent export (nndle), with three contamination values and from 12 fractionation experiments replicates merged (see methods). **D** Summary sketch: mRNAs with long poly(A) tail are synthesized and rapidly deadenylated in the nucleus. mRNAs with shortened tails are exported to the cytoplasm where they are translated, until progressive deadenylation leaves mRNAs without poly(A) tails, which are decapped and degraded.

## Discussion

Active mRNA degradation is a key mechanism of post-transcriptional gene regulation. Complete deadenylation of poly(A) tails is typically required for decapping and degradation of transcripts. The rates of deadenylation are gene-specific and therefore central determinants of RNA stability (Brown & Sachs, 1998; Cao & Parker, 2001; Eisen et al., 2020). Biochemical and labeling studies indicate the synthesis of poly(A) tails of ca. 250 nt by nuclear poly(A) polymerase (Eckmann et al., 2011), which are then exported and deadenylated in the cytoplasm (Brawerman & Diez, 1975; Sheiness & Darnell, 1973), although a genome-wide study on deadenylation rates concluded that mRNAs may emerge in the cytoplasm with poly(A) tails of ca. 130-150 nt on average (Eisen et al., 2020).

We found that poly(A) tails of nascent transcripts, which we identified from unspliced reads in FLAM-seq data, support the hypothesis of genome-wide synthesis of long poly(A) tails with a tail length of 200-250 nucleotides, consistent across samples and conditions. We validated the detection of unspliced reads by performing splicing inhibition and biochemical fractionation experiments *in vivo* and *in vitro*, showing that unspliced transcripts are enriched in the nucleus and increased upon splicing inhibition. We further analyzed published Nanopore direct RNA sequencing datasets of highly pure nascent, chromatin-associated transcripts (Drexler et al., 2020) and confirmed genome-wide synthesis of long poly(A) tails. A previous study applying Nanopore long-read sequencing on RNA from a human cell line also showed that reads containing intronic sequences had longer poly(A) tails, but these were attributed to possibly aberrant intron-retaining transcripts (Workman et al., 2019). Our computational pipeline relied on identification of unspliced and polyadenylated transcripts which are likely products of RNAs undergoing post-transcriptional (terminal) intron splicing. The extent of post-transcriptional versus co-transcriptional splicing is a matter of ongoing research. Recent studies indicate the occurrence of post-transcriptional terminal intron splicing for up to 50% of all mammalian genes (Coté et al., 2020; Drexler et al., 2020).

To rule out that our analysis was limited to specific subsets of transcripts, which could have non-stereotypical splicing patterns, we carefully quantified molecular properties such as intron length or expression levels of (a) transcripts with unspliced reads and (b) all transcripts detected. These analyses did not detect potential biases. However, we expect to miss unspliced RNAs because of limited read length, missing intron annotations or very long 3’UTR sequences.

Some unspliced transcripts had tails shorter than 150 nt in HeLa cells and we observed similar unspliced reads in cytoplasmic HeLa S3 fractions, which we attributed to transcripts with retained introns or erroneous annotations in the intron annotation we use for detection of unspliced reads.

Splicing inhibition affected bulk poly(A) length twofold: first, a mild global shortening was observed which could be explained by continuous deadenylation while mature RNA processing of newly synthesized transcripts is severely inhibited; second, an increase in transcripts with long poly(A) tails may have indicated compensatory mechanisms (e.g. for MYC, IER3, KLF5, genes with long 3’UTRs), or reflect nascent, unspliced RNAs not annotated by our computational pipeline for technical reasons (‘false negatives’). Stabilization of certain transcripts has been observed upon transcription inhibition (Shyu et al., 1989) and it is conceivable that similar mechanisms may sense blockage of splicing. PlaB has further been shown to induce cell cycle arrest and apoptosis, which may explain downregulation of the proto-oncogene MYC (Zhang et al., 2019). We further found poly(A) tail length of unspliced transcripts slightly increased upon splicing inhibition. Certain nuclear RNA decay pathways involve hyper-adenylation for clearance of aberrant transcripts by the nuclear exosome (Fasken & Corbett, 2009; Meola et al., 2016; Schmid & Jensen, 2018), and increased poly(A) length upon splicing inhibition may also indicate intermediates of nuclear decay.

Analysis of Nanopore direct RNA sequencing data (Drexler et al., 2020) extended our finding of global synthesis of long poly(A) tails beyond post-transcriptional splicing events, since non-polyadenylated nascent transcripts were tailed *in vitro* for Nanopore sequencing. 3’-end cleaved transcripts appeared either without or with long poly(A) tails. No evidence for the synthesis of intermediate poly(A) tail length significantly shorter than 250 nt was found in nascent RNA fractions. Poly(A) synthesis occurred rapidly, in the order of seconds (Elmar Wahle, 1995), which may explain emergence of bimodal length profiles at the poly(A) site in these datasets.

We did not find evidence for unspliced transcripts in nuclear fractions with poly(A) tails shorter than 100-150 nucleotides. This was also true upon splicing inhibition, which we expected to increase the dwell-time of unspliced mRNAs in the nucleus. We therefore hypothesize that completion of splicing could be required for shortening of tails.

Resolving the kinetics of the early phase of deadenylation using metabolic labeling and pulldown experiments revealed an increased poly(A) tail length compared to the steady-state poly(A) distribution. Surprisingly, the differences in poly(A) tail length for pulldown samples between labeling timepoints were minor for the majority of genes. For certain genes, such as IEGs or lncRNAs, a more pronounced shortening throughout the labeling period was observed. This result required for a rapid initial deadenylation step within minutes after synthesis, given the assumption of poly(A) tails being longer than 200 nt at the time of synthesis. We further applied SLAM-seq in combination with FLAM-seq which qualitatively resulted in a similar finding but suffered from technical challenges (Supplementary Discussion).

We observed significant shortening of tails in nuclear fractions of HeLa cells and mouse brain samples, which aligns with the hypothesis of rapid deadenylation. Our findings contradict earlier hypotheses describing export of non-deadenylated transcripts (Brawerman & Diez, 1975; Sheiness & Darnell, 1973), but support more recent models observing emergence of cytoplasmic tails with an average length of 130-150 nt (Eisen et al., 2020). A recent study investigating cytoplasmic and nuclear poly(A) tail length distributions in mouse 3T3 cell lines and ES cells also found and increased global nuclear tail length (Liu et al., 2021).

Nanopore direct RNA sequencing data of chromatin fractions together with 4sU enrichment for very short labeling periods of 8 min (Drexler et al., 2020) show that most poly(A) tails are still long without apparent nuclear shortening. Our chromatin fractionation experiments on the contrary show significant shortening of tails which can be explained either by a relatively long dwell time on chromatin upon which deadenylation occurs, which would be excluded by 4sU purification, or inefficient separation of nuclear and chromatin fractions. A similar argument relates to the 10-min. metabolic labeling performed in this study, where we observed significant shortening. We hence hypothesize that nuclear deadenylation may require a lag phase and/or release from chromatin, which could for instance involve a change in nuclear compartments.

We found evidence for gene-specific nuclear deadenylation since certain classes of genes show differences in tail length even after short labeling time points, when RNA is likely still localized in the nuclear compartment. Yet, differentiating the contributions of RNA export, nuclear deadenylation rates and nuclear decay on nuclear steady-state poly(A) distributions is in general very challenging through the lack of appropriate kinetic data.

An exception to nuclear deadenylation were lncRNAs, with more than 50% of nuclear lncRNAs not showing poly(A) tail shortening. Many lncRNAs function in the nucleus, for instance NEAT1 in paraspeckle formation (Fox et al., 2017), and further undergo distinct processing events compared to mRNAs, which may suggest coupling between nuclear deadenylation and other processing steps (Deveson et al., 2018; Mukherjee et al., 2017; Schlackow et al., 2017). In particular, we found a negative correlation between cytoplasmic-to-nuclear enrichment and poly(A) tail length in nuclear fractions, which lets us hypothesize nuclear deadenylation as a mechanism controlling RNA export which could be utilized to retain nuclear lncRNAs and/or to determine export rates via gene-specific deadenylation rates.

Quantitative modeling of deadenylation dynamics and mRNA steady state distributions predicts that nuclear deadenylation is by an order of magnitude faster than cytoplasmic deadenylation, which is expected given the significant shortening observed for chromatin and nucleoplasmic fractions within relatively short dwell times in the nucleus until RNA is exported. Interestingly, allowing poly(A) tail-length dependent export rates or nuclear mRNA decay dramatically improves the goodness-of-fit. Nuclear decay is mediated by the nuclear exosome which has a broad range with up to a 1000 mRNA targets reported in yeast (Delan-Forino et al., 2020), arguing for widespread nuclear decay which is consistent with our model. Similarly, poly(A) tails have been shown to increase export rates in *in vitro* experiments (Dower et al., 2004), although little is known about the impact of poly(A) tail length itself on export rates.

A number of deadenylase enzymes have been identified with CCR4-NOT and PAN2-PAN3 being most relevant for canonical deadenylation. A biphasic model has been proposed for yeast and mammalian deadenylation (Brown & Sachs, 1998; Yamashita et al., 2005) in which PAN2-PAN3 first trims long tails, and CCR4-NOT subsequently deadenylates tails until degradation. Whether this process is coordinated across nucleus and cytoplasm is not known, although both PAN2-PAN3 and CCR4-NOT are able to shuttle between cytoplasm and nucleus (Goldstrohm & Wickens, 2008) and have a divergent set of target genes in yeast (Tudek et al. 2021). A recent study investigating poly(A) tail dynamics of serum response genes after transcription induction identified the CNOT1 subunit of the CCR4-NOT complex as a nuclear deadenylase complex in mouse fibroblasts (Singhania et al., 2019). Whether the same enzyme complexes operate in different cellular compartments with different processivity is yet unknown.

In summary, we present evidence for rapid deadenylation of mRNAs occurring in the nucleus of mammalian cells (Fig. 4C). We propose that mRNAs are initially synthesized with a tail of 200-250 nt, which is then shortened in a splicing-dependent manner, with possible implications in the control of mRNA export. Similar to cytoplasmic deadenylation, nuclear poly(A) tail shortening is a gene-specific process which may serve as a nuclear control instance specifying the fate of mRNAs before they enter the cytoplasm.

## Authors contributions

Jonathan Alles: Study Design, Experiments, Computational Analysis, Manuscript Ivano Legnini: Study Design, Experiments, Computational Analysis Maddalena Pacelli: Mouse brain biochemical fractionation and FLAM-seq Nikolaus Rajewsky: Study Design & Supervision

## Acknowledgements

We thank C. Cerda Jara from the Rajewsky lab for preparation of mouse brain subcellular fractions and preliminary analysis, C. Quedenau from the BIMSB Genomics platform for PacBio Sequencing, S. Ayoub and M. Schott from the Rajewsky lab for help with library preparations, A. Rybak-Wolf, R. P. Alacaraz and A. Boltengagen from the BIMSB Organoids platform for help with mouse brain fractionations, N. Karaiskos and M. Jens from the Rajewsky lab, D. Schwabe from the Falcke lab and F. Silverii (GFZ) for helpful dicussions on modelling & data analysis, E. Conti and I. Schäfer (MPI Biochemistry) and E. Wahle (Halle U.) for helpful discussions and data interpretation. Illustrations in Fig. 2A, 3A, 3D and 4A were created with biorender.com. I.L. was supported by an EMBO long term fellowship (ALTF1235-2016). J.A. was supported by the MDC-NYU PhD exchange program. M.P. was supported by an Erasmus Mundus fellowship.

## Materials and Methods

### Experimental Methods

#### FLAM-seq library preparation and PacBio Sequencing

FLAM-seq library preparation was prepared according to (Legnini et al., 2019). In brief, polyadenylated RNA was isolated from total RNA using Illumina Truseq mRNA preparation kit. A GI tail was appended to selected mRNA using tailing reagents from USB poly(A) length assay kit. Tailed RNA was purified using 1.8x RNAClean beads. Tailed RNA was reverse transcribed using SMARTScribe Reverse Transcriptase Kit and isoTSO and RT primer 1 or 2. cDNA was purified using 0.6x Ampure XP beads and PCR amplified using PCR primer 1 or 2 and PCR Primer IIA using Advantage 2 DNA polymerase mix. FLAM-seq libraries were purified 2x using 0.6x Ampure XP beads.

PacBio adapters ligation and sequencing was performed according to standard procedures.

A detailed protocol for the FLAM-seq method can be found at https://protocolexchange.researchsquare.com/article/pex-398/v1

#### Splicing Inhibition using PlaB

Two 10 cm dishes of HeLa S3 cells per replicate cultured in DMEM medium were supplemented with 10 uL 100 nm Pladienolide B (PlaB) (dissolved in DMSO) or DMSO (control). Cells were incubated for 3h at 37°C and 5% CO2.

Cells were once washed with 5 mL cold PBS. 2 ml cold PBS, supplemented with 1:200 Ribolock and 1:100 Proteinase Inhibitor, was added to each plate and cells were scraped from dishes and collected in 2 mL tubes. Cells were spun for 5 min at 300 g and supernatant was removed.

For isolation of nuclei, the cell pellet was carefully resuspended in 500 uL 1x lysis buffer and incubated for 5 min on ice. A cushion of 500 uL 1x lysis buffer / 50% sucrose solution was pipetted under the cell lysate. Lysates were centrifuged through the cushion for 10 min at 16.000 g. The supernatant was removed and the nuclei pellet was again resuspended in 500 uL 1x lysis buffer and centrifuged through a sucrose cushion.

RNA from isolated nuclei was extracted using Trizol and Phenol-Chloroform extraction (Chomczynski & Sacchi, 1987). A DNAse cleanup step was added to remove gDNA, using Turbo DNA-free kit.

FLAM-seq libraries were prepared as described above.

### Predicting bivalent TSSs

HEK Flp-In T-rex cells cultured in four 15 cm dishes in DMEM medium. 4-Thiouridine (4sU, dissolved in DMSO) was added at a final concentration of 1 mM. Incubation was performed for 10, 15, 20, 45 and 90 minutes. For 0 min control, DMSO was added.

Cells were washed once with 10 mL cold PBS. 5 mL Trizol was added per dish and RNA was extracted using direct-zol RNA miniprep kit with DNAse gDNA removal.

Two biotinylation reactions (labeling buffer, 1mg/mL biotin-EZ-link) with each 100 ug total RNA input were prepared for each sample and incubated for 2h at RT in the dark.

Biotinylated RNA was purified using Phenol-Chloroform extraction. RNA was denatured for 3 min at 70°C and placed on ice.

Pulldown experiments were performed using MyOne Streptavidin C1 beads. 120 uL bead suspension were washed three times with 150 uL MPG buffer on a magnetic rack. Biotinylated RNA in 40 uL H2O was added to 120 uL MPG buffer on Streptavidin beads and incubated for 15 min at RT with rotation. Supernatants were separated from beads with bound biotinylated RNA by incubating on a magnetic rack for 1 min and collecting the supernatant as unbound fraction. Beads were washed three times with 150 uL pre-warmed MPG buffer (37°C). 150 uL 100 mM DTT was added and incubated for 5 min for elution of biotinylated RNAs as bound fractions. RNA from bound and unbound fractions were purified by Phenol-Chloroform extraction. FLAM-seq libraries were prepared from bound and unbound fractions as described above.

Dot blots were prepared by spotting 5 ug RNA from from each sample before and after biotinylation on a Amersham Hyperbond N+ membrane. The membrane was dried and crosslinked for with 2x 1200 uJ at 254 nm. The membrane was incubated with methylene blue for 10 min in order to record images of RNA quality per spot.

The membrane was blocked 20 min in Blocking Solution and probed for 10 min with 1:10.000 dilution of Strep-HRP antibody. Membranes were washed with each 2x for 5 min with 10%, 1% and 0.1% Blocking Solution. Signals of biotinylated RNA were recorded by chemiluminescence detection upon addition of ECL Selection reagent.

#### SLAM-Seq

HeLa S3 cells were cultured in DMEM medium. Cells were seeded on 6-well plates until reaching 70% confluency. For metabolic labeling the medium was supplemented with 500 uM 4sU and or DMSO control and incubated for 0 min, 90 min, 180 min. Cells were harvested and RNA was extracted using Trizol and Phenol-Chloroform extraction.

Polyadenylated RNA was extracted from 10 ug total RNA per sample using TruSeq RNA purification beads (Illumina) and eluted in 15 uL H2O.

GI tailing was performed using the USB poly(A) length kit (Thermo Fisher). 4 uL 5x tail buffer mix was added to 2 uL 10x tail enzyme mix and incubated for 1 h at 37°C. 1.5 uL Stop solution was added to quench the reaction. GI-tailed RNA was cleaned up using a 1.8x ratio of RNAClean XP beads. For alkylation reactions introducing T-C conversions, 15 uL GI-tailed RNA was incubated with 5 uL 100 mM iodoacetamide, 25 uL DMSO and 5 uL 500 mm NaPO4 pH 8.0 buffer for 15 min at 50°C. The reaction was quenched by addition of 1 uL 1 M DTT. RNA was purified using a 1.8x ratio of RNAClean XP beads. RNA was reverse transcribed by incubating 16 uL GI tailed RNA with 2 uL dC 3T UMI RT primer for 3 min at 72°C and placed on ice. 22 uL RT Mix (8 uL 5x RT buffer, 1.5 uL 100 mM DTT, 4 uL 10 mM dNTPs, 2 uL Ribolock, 2 uL IsoTSO 12 uMm, 2 uL SMARTScribe RTase, 2.5 uL H2O) were added and incubated for 1 h at 42°C, 10 min 70°C then 4°C hold. cDNA was purified using 0.6x ratio of AmpureXP beads. cDNA was eluted in 42 uL H2O. For PCR amplification 10 uL 10x Advantage 2SA PCR Buffer, 2 uL 10 mM dNTPs, 2 uL PCR Primer II A (12 uM), 2 uL Universal RV primer 10 uM and 2 uL 50x Advantage 2 Polymerase and 42 uL H2O were added and incubated using the following program 98°C 1 min [98°C 10 sec, 63°C 15 sec, 68°C 3 min] 68°C 3 min. cDNA libraries were cleaned up 2x 0.6x Ampure XP beads.

#### HeLa S3 Cell Fractionation

HeLa S3 cells were seeded in two 10 cm dishes per replicate in DMEM medium. Cells were washed once with cold PBS. 2 ml cold PBS, supplemented with 1:200 Ribolock and 1:100 Protease Inhibitor, was added and cells were scraped from dishes and collected in 2 mL tubes. 50 uL cell suspension was collected as input fraction.

The cell pellet was carefully resuspended in 500 uL 1x lysis buffer and incubated for 5 min on ice. A cushion of 500 uL 1x lysis buffer / 50% sucrose solution was pipetted under the cell lysate. Lysates were centrifuged into the cushion for 10 min at 16000 g. The supernatant was collected as cytoplasmic fraction. The nuclei pellet was again resuspended in 500 uL 1x lysis buffer and centrifuged through a sucrose cushion, the procedure was repeated once. Pellets were carefully resuspended in 100 uL Nuclear Buffer I. 1 mL Nuclear Buffer II was added, tubes were inverted 5 times and incubated for 15 min on ice. Suspensions were centrifuged for 10 min at 16.000 g. Supernatants were collected as nucleoplasmic fraction. Chromatin pellets were resuspended in 500 uL H2O. 50 uL of each fraction were collected for Western Blot analysis. 5 vol Trizol were added to resuspended fractions and RNA was extracted by Phenol-Chloroform extraction.

FLAM-seq libraries were prepared as described above.

For Western Blot analysis, 15 uL suspension from input, cytoplasmic and chromatin fraction, as well as 30 uL nucleoplasmic fraction were loaded on 12% SDS-PAGE gels and run according to standard protocols (Laemmli, 1970). Blotting was performed using BioRad Trans-Blot Turbo Transfer System and standard settings for 2 mini gels. Membranes were blocked using 5% skim milk in TBS-T buffer for 1h. Membranes were then probed with BCAP31, GAPDH and TBP-43 antibodies in 5% skim milk / TBS-T overnight. Membranes were washed 3x in TBS-T and incubated with 1:10.000 anti-mouse- or anti-rabbit-HRP antibodies in 5% skim milk / TBS-T buffer. Membranes were washed 3x with TBS-T buffer. Membranes were probed with ECL Select Solution and imaged.

For certain experiments membranes were stripped and re-probed with antibodies.

#### Mouse Brain Cell Fractionation

The method for separation of nuclei and cytoplasmic fractions was adapted from Sigma-Aldrich single nuclei isolation protocol (nuclei EZ prep nuclei isolation kit, Product number NUC-101). The mouse brain tissue was thawed in a Petri dish and 1 mL EZ lysis buffer was added. A razor blade was used to cut the brain into two hemispheres, which each were used as replicates. The tissue was transferred in cold Dounce homogenizer and chopped with 20-25 strokes with pester A and 15 strokes with pester B. Homogenate was transferred into a 15ml Falcon tube with wide-bore pipette. An aliquot was saved for RNA quality control. After addition of 2 mL of EZ-lysis buffer, the falcon was vortexed briefly and the sample was incubated 5 min on ice. An aliquot of the whole cell lysate was saved for Western Blot. The tube was centrifuged at 500 g for 5 min at 4°C and supernatant was collected as cytoplasmic fraction. An aliquot was saved for Western Blot analysis. The nuclei pellet was resuspended in 4 mL EZ lysis buffer, incubated 5 min on ice and centrifuged again at 500 g for 5 min at 4°C. The supernatant was carefully aspired and the pellet was resuspended in 200 ul PBS-BSA 0.01%. An aliquot of the nuclei fractions was saved for Western Blot analysis. 10 uL nuclei sample was stained with DAPI to assess the quality of the nuclei suspension. All steps were performed on ice to minimize nuclei damage and maintain RNA quality. Western blot analysis of mouse brain fractions was performed as described above. RNA was extracted from fractions by Phenol-Chloroform extraction. FLAM-seq libraries were prepared as described above.

#### Computational Methods Processing of FLAM-seq datasets

FLAM-seq datasets were processed using the FLAMAnalysis pipeline (https://github.com/rajewsky-lab/FLAMAnalysis) to identify gene identities as well as poly(A) tail length and tail sequence for each read. For analysis of 3’UTR isoforms, datasets were processed as previously described (Legnini et al., 2019) in order to group reads by 3’UTR isoforms. For human datasets, reads were mapped against human genome reference hg38 and human Gencode annotation version 28, both complemented with sequences from Thermo Fisher ERCC spike ins. For mouse datasets, reads were mapped against mouse GRCm38 and mouse Gencode annotation GRCm38.101.

#### Identification and analysis of unspliced reads

For identification of unspliced reads from FLAM-seq datasets, reads were required to overlap with a curated set of intron coordinates as well as 3’UTRs in order to exclude events of partial splicing or intron retention.

Intron sets were curated as follows. We first downloaded all introns, exons and 3’UTR annotations from UCSC table browser (hg38, Gencode v24 for human, mm10, Gencove VM23 for mouse), then filtered for only protein coding genes, for which we presumed a better annotation, and filtered out those overlapping between each other with bedtools intersect (Quinlan & Hall, 2010). Along the same line, we filtered out all introns overlapping with annotated exons, also with bedtools intersect, not allowing any overlap.

Intronic reads were filtered using `bedtools intersect -a bam_alignment -b intron.bed`, filtering out overlaps shorter than 50 nt and retaining only unspliced reads, and restricted to those reads overlapping annotated 3’UTRs, in order to filter out potential artifacts such as internally primed reads, through `bedtools intersect -a intronic_reads.bed -b 3utrs.bed`.

For visualization of intronic poly(A) tail length distributions, intronic read counts were normalized to absolute reads counts for each dataset.

For downsampling analysis, intronic reads and detected genes in HeLa S3, iPS and Organoid datasets were merged. Intronic reads were downsampled and for each downsampled read set the number of detected genes as a fraction of total genes in the merged dataset was calculated.

To investigate, for how many detected genes in FLAM-seq datasets identification of unspliced reads is possible, coordinates of introns in curated intron set were compared to gencode annotation of genes using `bedtools intersect -s -wa curated_introns.bed -wb gencode_genes.bed.`

Intersected coordinates were then filtered for genes where the intron end is < 3 kbp away from the annotated gene end, to get a realistic estimate of detectable genes since >95% of all reads are shorter than 3 kb.

In search of potential biases in our analysis, we compared the length distribution of introns to which unspliced reads were mapped to all introns (Fig. S1F). We also compared the distribution of intron length derived from Gencode annotation (Frankish et al. 2021) for genes with detected unspliced transcripts against all genes per sample (Fig. S1G).

For investigation of gene expression dependent changes in poly(A) tail length genes from merged datasets for HeLa S3, iPS and Organoid datasets were binned by gene counts. Intronic reads with respective poly(A) tail lengths were assigned to genes in each expression bin (Fig. S1H-I).

#### Analysis of Nano-Cop nascent RNA Nanopore sequencing

We analyze previously published Nanopore direct RNA sequencing datasets (Drexler et al., 2020) to investigate poly(A) tail length of nascent, chromatin associated RNA. Fastq records and processed data were downloaded from GEO GSE123191. Fastq reads were mapped using Minimap2 (version 2.16-r922) (Li, 2018) for direct RNA sequencing `minimap2 -ax splice -uf -k14 -t 8 index fastq > sam`

Aligned reads were annotated using featureCounts and human gencode version 28 .gtf annotation `featureCounts -L -g gene_name -s 0 -t gene -O --fracOverlap 0.3 -R CORE -a gtf -o out sam` Reads were grouped by provided read end annotation defining how read ends overlap with gene features (i.e. intronic, poly(A) site). Reads were annotated by provided poly(A) tail length estimates. For extraction of unspliced reads, we applied our computation pipeline as described above.

#### Browser visualization of FLAM-seq alignments

Visualization of FLAM-seq alignments including poly(A) tails on genome browsers was achieved by appending poly(A) tail sequences to the alignments, and loading these new alignments on IGV (or Gviz, or UCSC genome browser), taking advantage of the fact that these browsers mark mismatches with a different color for each nucleotide (thus coloring mismatched Ts, from the complementary DNA of the poly(A) tail, with a different color than the bulk of the aligned read). Briefly, the poly(A) tail sequences are retrieved from the output of the FLAMAnalysis pipeline (the *cleaned_tail_lengths.txt* file) and appended to the corresponding read of the bam file, also produced by the FLAMAnalysis pipeline. The CIGAR string is modified accordingly, appending a stretch of mismatches with the same length as the appended poly(A) tail.

#### Analysis of intron length

For comparison of intron length between genes with unspliced reads and all genes in each dataset, intron lengths of all annotated transcript isoforms for each gene were extracted from Homo Sapiens Gencode GTF annotation v28. Intron distributions for genes with detected unspliced reads were compared to the background of all detected genes in each dataset.

For comparing length of non-exon overlapping intron sequences (used for identification of unspliced reads, s. above), length distribution of introns overlapping with unspliced reads were compared to all introns in the curated intron set.

#### Analysis of half-life / 3’UTR length for PlaB splicing inhibition

For analysis of gene half-lifes between genes with altered median poly(A) tail length upon splicing inhibition, genes were first grouped into bins based on differences in median poly(A) tail length between PlaB treated and control samples. Genes with differences >50 nt were grouped in “shorter poly(A) PlaB” or “longer poly(A) PlaB” bins while remaining genes were classified as “unchanged poly(A) PlaB”. Control datasets comprised random genes of identical sample site as “shorter”/”longer” bins.

RNA half-life estimated per gene for HeLa cell lines were taken from Tani et al., 2012 and plotted for each bin.

3’UTR isoforms for reads from FLAM-seq datasets were annotated as previously described (Legnini et al., 2019) and 3’UTR length was plotted for each bin.

### Metabolic labeling experiments

4sU metabolic labeling and pulldown experiments were analyzed by first quantifying poly(A) tail length for each read using the FLAMAnalysis pipeline. Intronic reads were extracted as described above. Poly(A) tail lengths between pulldown and supernatants were compared by calculating the differences in median poly(A) tail length of genes in pulldown datasets with the merged supernatant dataset for genes with more than 3 counts.

For analysis of SLAM-Seq datasets, reads were first processed applying the FLAM-seq pipeline to retain only reads with valid poly(A) tails. In a next step, filtered alignment files (*_cleaned.bam) were annotated with an MD tag using `samtools calmd` (Li et al., 2009) with hg38 reference genome used for FLAMAnalysis pipeline. To exclude low quality reads with aberrant mutation patterns reads with >30 overall mutations and read quality <85 were excluded. We clipped the first 20 bases of each read as we observed an enrichment of mutations towards the read start which reflect mapping artefacts.

As proposed by (Jürges et al., 2018) we estimate the ‘labeling rate’ plabel of T-C conversions from the fraction of observed T-C mutations over all bases sequences in a given dataset. We estimate the background (T-C) ‘error’ rate perror as the average mutation rate for all non-T-C mutations. For each read estimate the likelihood of an observed number of T-C mutations under a ‘labeling’ model versus an ‘error’ model. We model the probability of observing k T-C mutations in a read containing n T’s under each model as a binomial with B(n;k;plabel) or B(n;k;perror). We set a cutoff for flagging each read as ‘labeled’ if the log likelihood >1.15, which minimizes the number of false positives in datasets without 4sU labeling. We then report a matrix with labeled and unlabeled reads for each gene.

Median poly(A) tail length per gene was compared between labelled reads and all reads per gene in each dataset by subtracting the median for genes with detected labeled reads and more than 3 counts in total.

### Analysis of biochemical fractionation data

FLAM-seq datasets from biochemical fractionation experiments were processed using the FLAMAnalysis pipeline. 3’UTR isoforms for reads mapped to a given gene were annotated as described previously (Legnini et al., 2019).

Intronic reads were identified for each dataset as described above.

For comparisons of all poly(A) tail length profiles between fractions, poly(A) tail length densities were calculated for each replicate and fraction for poly(A) tail length bins of 10 nt. Mean and standard deviations were calculated for each poly(A) bin and plotted. Average density distributions for intronic reads calculated similarly and smoothed using R ‘smooth’ function.

To investigate sources of variation for poly(A) tail length profiles between different biological replicates we first calculated a scaling factor for each replicate as average scaled median poly(A) length for each fraction of a replicate.

We apply a linear model to predict poly(A) tail length in a fraction of the variables ‘scaling factor’, ‘RNA concentration’, ‘fraction intronic reads‘ and ‘cellular compartment’ and evaluate model performance using different predictors to identify factors best explaining variability in poly(A) tail length distributions.

Scaled poly(A) tail length distributions were calculated by dividing poly(A) tail length for each read by the scaling factor the fraction.

For analysis of poly(A) tail length of genes with or without associated intronic reads, bulk reads were split by genes in each bin for each replicate and average densities were calculated as above.

Analysis of gene counts by poly(A) tail length bin was performed by binning median poly(A) tail length per gene and aggregating gene counts for genes in each bin. For analysis of intronic poly(A) tail length by poly(A) bin, genes were grouped by median poly(A) length bin and poly(A) tail length of intronic reads for genes in each bin were aggregated.

For analysis of gene half-life by poly(A) tail bin, half-life data of HeLa cell lines were downloaded from (Tani et al., 2012) and plotted for each gene in poly(A) tail length bins.

3’UTR isoforms for genes with distal and proximal isoforms were extracted from 3’UTR annotation matrices. 3’UTR length was plotted for proximal and distal isoforms on poly(A) tail length bins for isoforms.

lncRNA genes were extracted based on bioMart (Durinck et al., 2005) gene biotype annotation.

For comparison of median poly(A) tail length per gene between fractions, genes were filtered to have more than 5 counts in each fraction. Cytoplasmic-to-nuclear ratios for each gene were taken from processed ENCODE data (Yi et al., 2018) and plotted again median poly(A) tail length in each fraction for genes with > 25 counts.

All above analyses were performed on a merged dataset of all replicates for each fraction.

### Mathematical modelling of nuclear and cytoplasmic mRNA tail lengths distributions

In order to understand whether our hypothesis of a rapid nuclear deadenylation process is consistent with the observed tail length distributions at synthesis and at steady state, we conceived a simple model describing the amount of mRNA in a cell with a given tail length as a function of transcription, export, deadenylation and decay. For simplicity, we model the total amount of mRNA in the nucleus and in the cytosol as a function of global rates, thus not estimating gene-specific parameters as for example in Eisen et al. (2020), where much deeper sequencing data were produced for both steady state and metabolic labeling experiments.

In this model, we assume that a single cell contains 200,000 mRNA molecules at steady state, sorted between nucleoplasm, chromatin and cytosol according to certain ratios estimated by experimental quantification of total RNA masses obtained from 12 fractionation experiments (assuming the total amount of RNA in the fractions reflects the amount of mRNA, see Table S1). We use these ratios to compute the amount of mRNA molecules in the nucleus and in the cytosol, at the steady state, for each tail length between 1 and 501 nt, directly from FLAM-seq data. To account for variable contamination of the nuclear fraction with cytoplasmic mRNA, we scaled these distributions according to a contamination of 0, 25 and 50%. We then simulate the amount of nuclear and cytosolic mRNA for each tail length, as resulting from the steady state solutions:

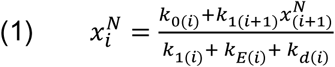

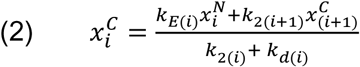

of the ODEs:

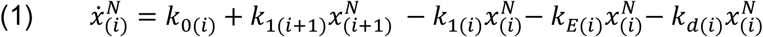

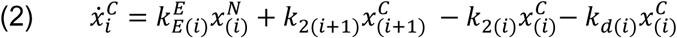

The amount of nuclear mRNA *x^N^* with tail length *i* is a function of a transcription rate k_0_ (the only zeroth-order process in the chain), a deadenylation rate k_1_ and the amount of mRNA with tail length *i* +1, an export rate kE, and in one case, to which we will come, a decay rate k_d_. Conversely, the amount of cytoplasmic mRNA *x^C^* with tail length *i* is a function of the export rate and the amount of nuclear mRNA with tail length *i*, a cytoplasmic deadenylation rate k_2_ and the amount of cytoplasmic mRNA with tail length *i* +1, and a decay rate kd. Some of these rates are tail-length dependent. k_0_, the transcription rate, is the product of a synthesis rate a and a synthesis tail length probability function (*slp*). The decay rate is computed as a logistic function of parameters ad, md and vd, as previously done in Eisen et al. (2020), since it describes the fact that tails are shortened to a certain length before decay by decapping and exonucleolytic activities can proceed. Nuclear and cytoplasmic deadenylation rates are not tail length-dependent. For the export rate kE, we allow two possibilities. We first set it as a constant (i.e. not depending on tail length) and we observed that the model would produce a large buildup of completely deadenylated molecules in the nucleus, which does not reflect common knowledge and observations. To account for this, we either allowed decay of short-tailed mRNAs in the nucleus (“nuclear decay – constant export” or ndce), or we added a second term to the export rate, modelled as a logistic of parameters aE, mE and vE, which would reflect a second scenario where mRNAs with a shortened tail are exported more efficiently (“no nuclear decay – logistic export” or nndle). Both these models were able to reproduce the experimentally determined poly(A) tail profiles, cutting down the residuals with the data of one order of magnitude with respect to the basic model with no nuclear decay and constant export.

With these two models, we proceeded to estimate the parameters by optimization. We used the sum of the squared differences between the simulated and measured nuclear and cytoplasmic mRNA amounts as a cost function to minimize. For minimizing it, we used the limited memory, box-constrained version of the Broyden–Fletcher–Goldfarb–Shanno method (L-BFGS-B) available within the optimx R package. We first make a guess of the initial parameters as explained below, then sample 1,000 times from a normal distribution built around each of these initial guesses with a standard deviation of 50% (and limited by generous lower and upper limits: if the sampling exceeds the limits, there are substituted with the initial guess), then run the optimization function to estimate a distribution for each parameter. This process is repeated taking into account three different contamination rates of FLAM-seq nuclear data, and for the two proposed models allowing nuclear decay and keeping export constant, or not allowing nuclear decay and adding a logistic export term. An example for the two models, along with that of the basic model which does not allow nuclear decay nor tail-dependent export, is shown in fig. 4B. We tested the models’ sensitivity to the parameters and observed that it behaved as expected (for example, increasing the synthesis rate increases the amount of mRNA in both nucleus and cytosol, increasing export rate increases at amount of mRNA in the cytosol at the expense of the nucleus etc.). Interestingly, given the tail dependency of decay, we observed that the cytoplasmic steady state mRNA level is strongly dependent on both nuclear and cytoplasmic deadenylation rates. In fig. S4E, we show as an example that doubling or halving the nuclear deadenylation rate results in changes of cytoplasmic mRNA level.

The parameters to initialize the optimization were estimated as follows. The slp is estimated with a loess fit of all intronic reads from nucleoplasm and chromatin of 12 fractionation experiments, and peaks at ca. 215 nt (Fig. S6A). A global synthesis rate is computed as the sum of all synthesis rates of all genes from Eisen et al., (2020), rescaled according to gene expression in our dataset (i.e. we kept only genes which were expressed in our data, and increased the total transcription rate by the ratio between the expression of these genes and all genes in our data). The global transcription rate was further scaled to estimate how many mRNA molecules are produced every minute in a cell containing 200,000 molecules at the steady state (Fig. S6A). A similar estimate (640 mol/min as compared to 712 mol/min) was obtained from a completely independent project carried on in our lab (Haiyue Liu, manuscript in preparation), where single-cell RNA-sequencing of cells labeled with 4-thiouridine was used to estimate transcription rates in single cells. The deadenylation rates k_1_ and k_2_ were set equal, according to the distribution of deadenylation rates from Eisen et al., (2020), weighted for gene expression in our data (Fig. S6A). The decay rate parameters were also set identical to Eisen et al., (2020), with the ad expressing the rate itself also weighted for gene expression in our data (fig. S6A). For the export rate, we set it around 0.02 min^-1^, estimating a nuclear half-life of ca. 30 min and consistent with metabolic labeling-based measurements in Drosophila by Chen and van Steensel (2017). For the logistic export rate, we adjusted the aE, mE and vE parameters from initial optimizations to 1.2, 5 and 10. These values are somehow arbitrary and are introduced, as previously stated, to eliminate the buildup of deadenylated nuclear molecules which happens if no nuclear turnover or no length-dependent export is allowed. As previously described, we sample 1,000 times from the distributions built around the initial guesses, and use these samples to initialize optimization with the merged FLAM-seq data from 12 fractionation experiments, with the two ndce and nndle models and with three values of contamination. We then retain the iterations with successful convergence and compute the parameters distributions shown in fig. S6B and C. The merged estimates from the two models and the three contaminations were used for the distribution shown in Fig. 4C. With both models and only 0.25 contamination, we separately repeated the optimization for the 12 individual fractionation experiments and show the means in fig. S6D.

### Supplementary Discussion

SLAM-seq is a fast and robust protocol for investigating RNA dynamics over time by labeling newly synthesized RNA, which can be identified in high-throughput sequencing experiments through introduction of T-C conversions in labeled RNA (Herzog et al., 2017). We combine the SLAM-Seq approach with our FLAM-seq method to simultaneously query poly(A) tail length of labeled reads.

Newly synthesized RNA incorporates 4sU which is added to the cell medium. 4sU moieties within RNA are alkylated by iodoacetamide (IAA) causing T-C mismatches during cDNA synthesis at the positions of 4sU incorporation. We perform 4sU labeling in HeLa S3 cell lines and subsequent poly(A) selection and GI tailing before IAA alkylation. Alkylation reactions are performed at basic pH and 50°C which may cause hydrolysis. While this has been carefully investigated for quantification of gene expression (Herzog et al., 2017), it is unclear whether poly(A) length quantification could be affected by harsh reaction conditions. To minimize the effects of possible hydrolysis on poly(A) tails, we adapted the FLAM-seq protocol such that we append a GI tail to RNA poly(A) tails before alkylation. RNA fragmentation in the tail would cause loss of the GI tail and exclude those RNAs from being reverse transcribed using the FLAM-seq poly(C) RT primer.

We performed labeling of HeLa S3 cell lines for 0 (control), 90 and 180 minutes. We developed a computational pipeline to extract mutations with respect to a reference genome and applied a likelihood model similar to the GRAND-SLAM approach (Jürges et al., 2018) to categorize reads as newly synthesized (“labeled”) or pre-existing (“unlabeled”) based on observed T-C mutations over a background model of expected T-C mutations estimated from all mutations in a dataset.

We observed an increasing number of average T-C mutations per read proportional to the 4sU labeling time. The average number of total mutations per read increases similarly, yet we observed more non-T-C mutations for longer labeling (Fig. S3I). Calculating the distribution of mutation events over all possible nucleotide conversions showed the expected enrichment of T-C conversions for 90 min and 180 min labeling while non-T-C mutations occurred at similar frequencies (Fig. S3J). Overall, we categorized 15% of all reads as labeled after 90 min 4sU and ca. 22 % after 180 min labeling. The fraction of labeled reads appeared slightly higher than the fraction of labeled total RNA in pulldown experiments (10% for 90 min labeling) and may be influenced by differences in labeling between ribosomal RNA, which makes up the majority of signal in pulldowns, and mRNA in SLAM-Seq.

Less than 1% of reads were flagged as “labeled” in 0 min control experiments, indicating high specificity of our computational approach (Fig. S3K).

Comparing poly(A) tail length distributions between labeled and total reads from SLAM-Seq experiments showed a modest increase in poly(A) tail length of newly synthesized RNAs as observed in HEK pulldown experiments (Fig. S3H).

Unspliced reads had longer poly(A) tails, yet with a significant number of reads having tails shorter than 100 nt as observed for bulk HeLa S3 datasets (Fig. S3L)

Poly(A) tail length distributions of mitochondrial transcripts also peaked at 50 nt both for labeled and total reads categories (Fig. S3M). Comparing the median poly(A) length for each gene between labeled and all reads showed a median increase in poly(A) length for labeled reads of 10 to 15 nt for 90 min and 180 min labeling respectively (Fig. S3K). We did note an overall shift in global poly(A) tail length profiles between individual timepoints, which is unexpected since the SLAM-Seq protocol should preserve the steady-state. 0 min and 90 min labeling samples had longer tails (median 102-106 nt) than 180 min labeling (median 66 nt), which we attributed to variability introduced by the experimental setup which required extensive handling of ultra-low RNA quantities.

We conclude that combining the SLAM-Seq method with full-length RNA sequencing enables parallel investigation of RNA dynamics and poly(A) tail length but will require further optimization of the experimental protocol to avoid batch effects between labeling timepoints.

### Data availability

Raw and processed FLAM-seq datasets related to this study can be accessed at Gene Expression Omnibus (GEO) under GSE188539.

---

**Supplement Fig. 1.**
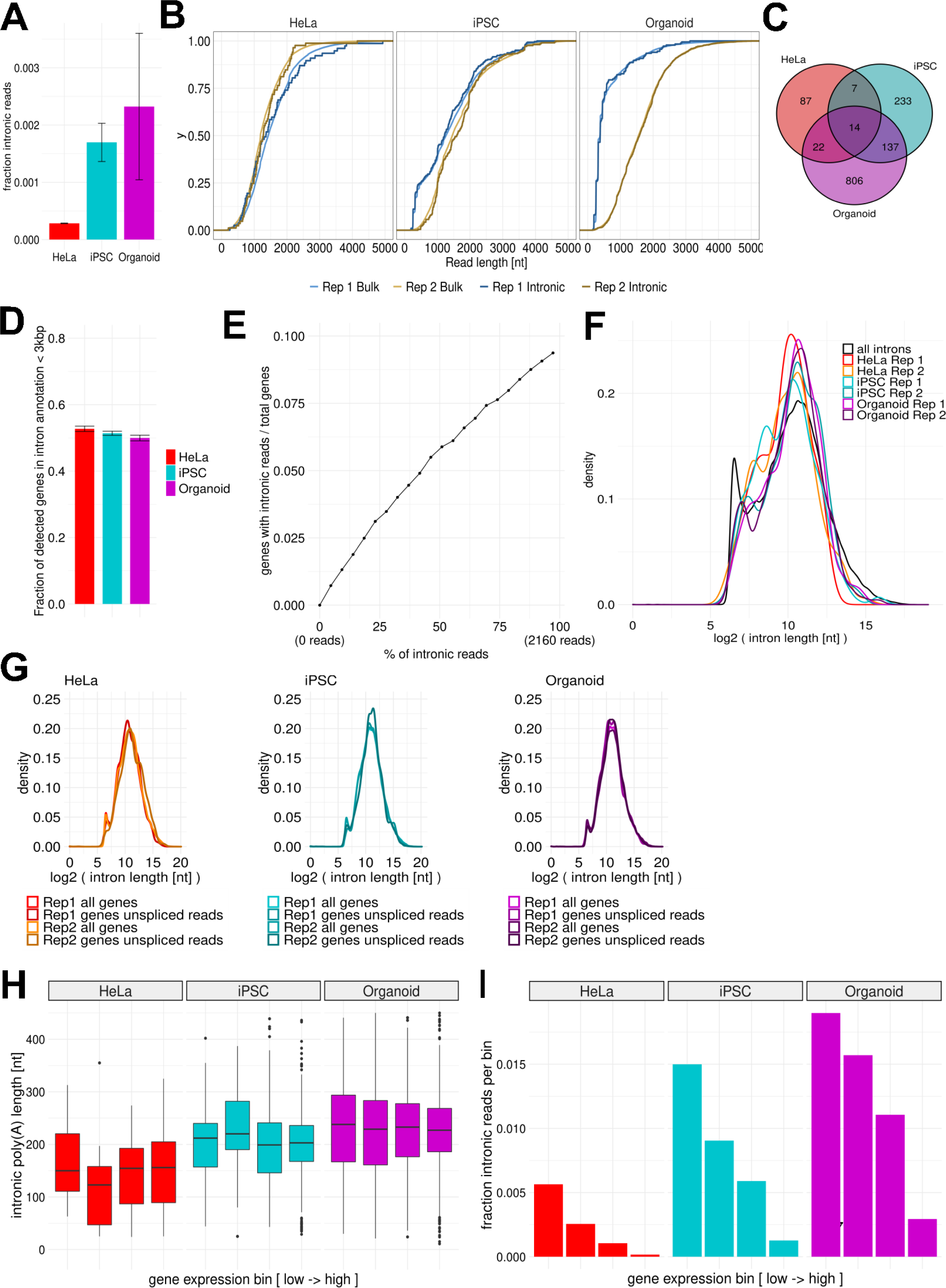
Unspliced RNAs have long poly(A) tails. **A** Fraction of unspliced, intronic reads in HeLa S3, iPSC, Organoids FLAM-seq datasets. Error bars indicate standard error of the mean for 2 replicates. **B** Cumulative distributions of raw read length for intronic and bulk reads for HeLa S3, iPSC and Organoid datasets. **C** Venn diagram of genes with detected unspliced (intronic) reads per dataset. **D** Fraction of detected genes in FLAM-seq HeLa S3, iPSC, and Organoids dataset represented in the intron annotation database a maximum intron distance of 3kb from gene end. **E** Downsampling analysis: Intronic reads were downsampled to respective fraction of total reads and the fraction of genes with intronic reads was compared to the total of detected genes. **F** Intron length of unspliced introns compared to distribution of all annotated introns for identification of unspliced molecules. **G** Intron length distribution from Gencode annotation comparing all genes detected in a dataset with those genes with associated unspliced reads. **H** Binning of intronic reads poly(A) length distributions by gene expression for merged datasets. Number of reads per bin is indicated below the boxplot. **I** Fraction of intronic reads by gene expression bin for merged datasets.

**Supplement Fig. 2.**
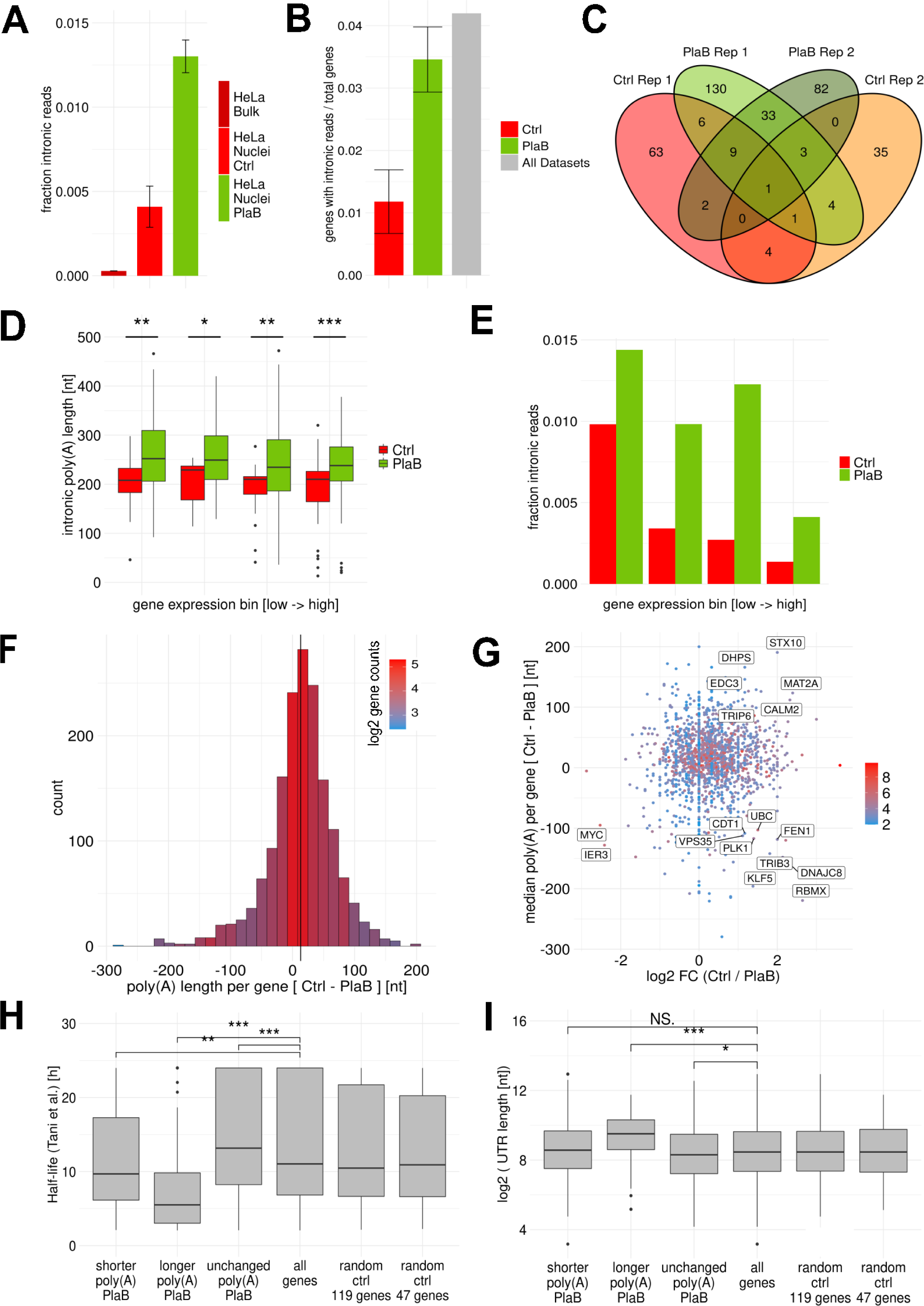
Unspliced RNAs have long poly(A) tails. **A** Fraction of intronic reads for HeLa Bulk, Nuclei Ctrl and Nuclei PlaB conditions. Error bars denote standard deviation of 2 replicates. **B** Fraction of genes with detected unspliced, intronic reads normalized to total number of genes detected in each dataset. Error bars denote standard deviation of 2 replicates. **C** Venn diagram of genes with detected unspliced (intronic) reads per dataset. **D** Binning of intronic reads poly(A) length distributions by gene expression for merged datasets. Number of reads per bin is indicated below the boxplot. Asterisks indicate significance in difference between Ctrl and PlaB (two-sided t-Test). **E** Fraction of intronic reads by gene expression bin for merged datasets. **F** Median poly(A) length difference per gene comparing HeLa S3 nuclei control versus PlaB treatment. Color indicates average log2 gene counts. Vertical line indicates mean. **G** Difference in median poly(A) length per gene between PlaB and control conditions compared to fold change in gene expression. Highlighted are genes with most prominent differences. **H** Distributions of gene HeLa S3 half-life (Tani et al.) for bins of genes with change in poly(A) length upon PlaB splicing inhibition and control distributions. **I** Distributions of transcript 3’UTR length for bins of genes with respective change in poly(A) length upon PlaB splicing inhibition and control distributions.

**Supplement Fig. 3.**
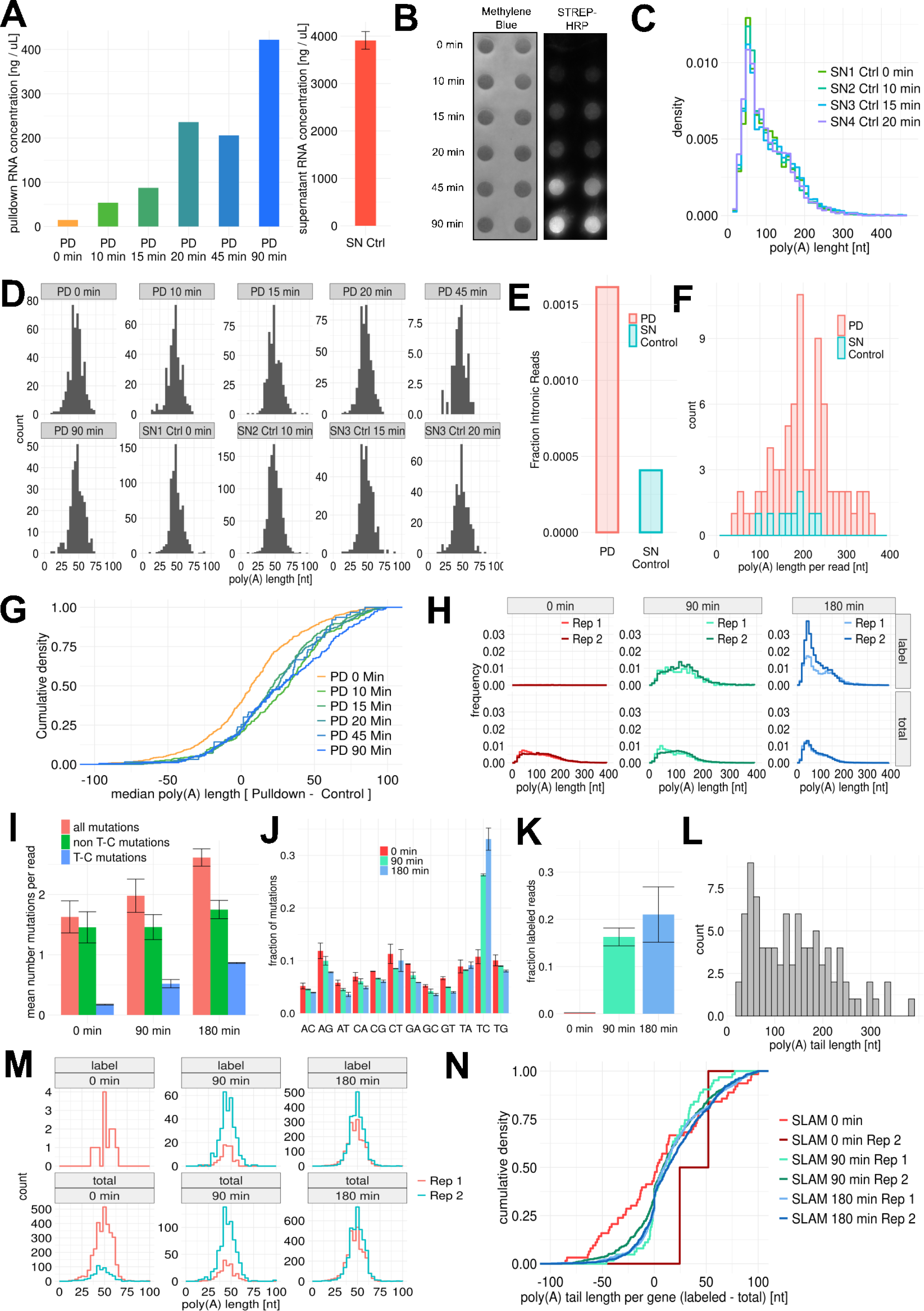
Metabolic labeling indicates rapid shortening of poly(A) tails after synthesis. **A** RNA concentrations obtained from pulldown (PD) fractions after indicated time points. **B** Dot blot with Methylene-Blue staining and Strep-HRP signal against biotinylated RNA before pulldown. **C** Poly(A) length distribution of supernatant (SN) fractions for 0 min to 20 min labeling timepoints. **D** Poly(A) length distributions of mitochondrial transcripts in PD and SN fractions. **E** Fraction of intronic reads in PD and SN fractions. **F** Poly(A) tail length distributions of intronic reads from PD and SN fractions. **G** Cumulative density distribution median poly(A) tail length difference per gene between PD and SN fractions. **H** poly(A) tail length profiles of labeled reads and all reads in SLAM-Seq experiments for 0 min, 90 min, and 180 min labeling. **I** Average number of mutations per read in SLAM-Seq datasets considering all mismatches per read, T-to-C mismatches or non-T-C mismatches. Error bars denote standard deviation of 2 replicates. **J** Observed nucleotide conversions in SLAM-Seq datasets normalized to all possible conversions. Error bars denote standard deviation of 2 replicates. **K** Fraction of reads assigned as ‘labeled’ in SLAM-Seq datasets. Error bars denote standard deviation of 2 replicates. **L** Poly(A) tail length distribution of intronic reads in merged SLAM-Seq datasets. **M** Poly(A) tail length distributions of mitochondrial transcripts in labeled or total poly(A) tail length bin. **N** Cumulative density distribution of median poly(A) tail length difference per gene between SLAM-Seq total and labeled reads.

**Supplementary Fig. 4.**
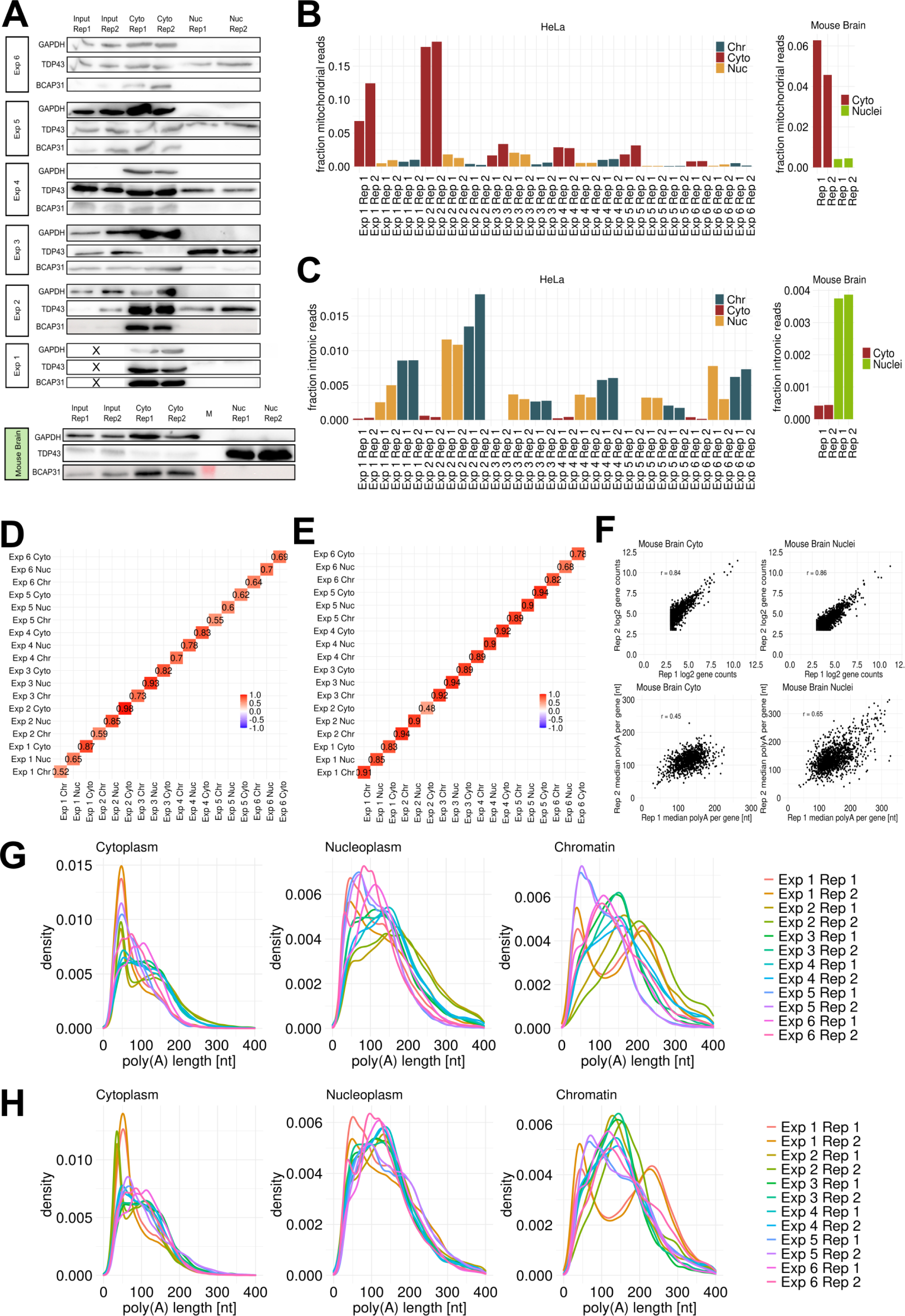
Biochemical fractionation reveals nuclear deadenylation. **A** Western Blot analysis of fractions obtained from biochemical fractionation of HeLa S3 cell lines and mouse brain samples. Markers used are GAPDH as cytoplasmic marker, TDP43 as cytoplasmic and nuclear marker and BCAP31 as ER marker. **B** Fraction of mitochondrial reads in each replicate for cytoplasmic, nucleoplasmic and chromatin fractions. Left: HeLa S3 cell lines Right: Mouse brain. **C** Fraction of intronic reads in each replicate for cytoplasmic, nucleoplasmic and chromatin fractions. Left HeLa S3 cell lines Right Mouse brain. **D** Correlation coefficients for median poly(A) tail length per gene between HeLa S3 technical replicates and corresponding fractions for genes with more than 10 counts. **E** Correlation coefficients for gene expression counts between HeLa S3 technical replicates and corresponding fractions for genes with more than 10 counts. **F** Comparison of gene expression counts per gene (top) and median poly(A) tail length per gene (bottom) for HeLa S3 mouse brain samples for genes with more than 8 counts. **G** Poly(A) tail length densities for HeLa S3 samples by fraction. **H** Poly(A) tail length densities for HeLa S3 samples by fraction scaled by average poly(A) tail length of sample over all fractions.

**Supplementary Fig. 5.**
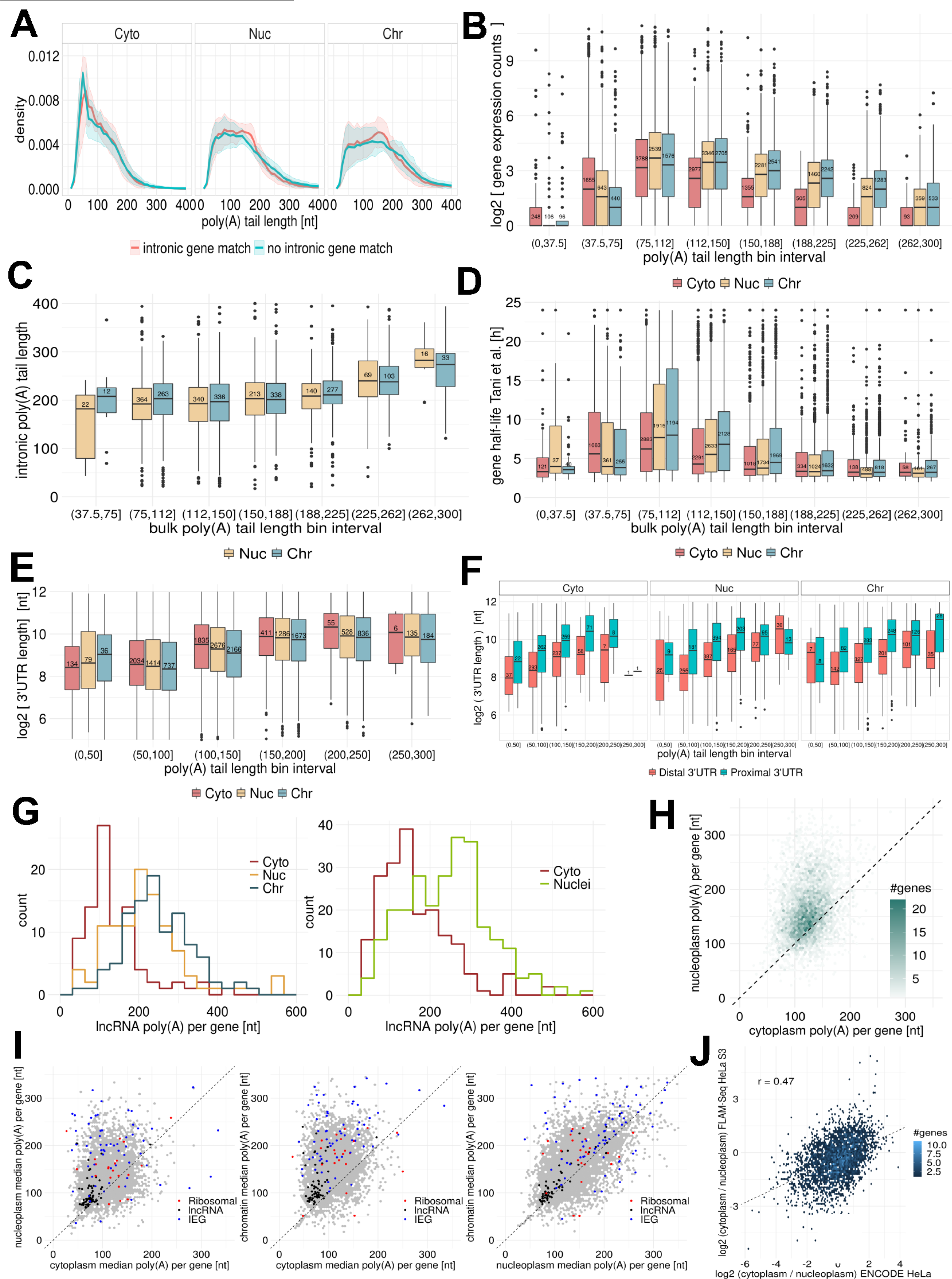
Biochemical fractionation reveals nuclear deadenylation. **A** Bulk poly(A) tail length distributions of genes with detected intronic reads (‘intronic gene match’) versus genes without detected intronic reads for HeLa S3 cell fractions. **B** Molecule counts for genes binned by average poly(A) tail length per gene. Numbers indicate the number of genes in each bin. **C** Poly(A) tail length of intronic reads binned by median poly(A) tail length of associated genes. Numbers indicate the number of intronic reads in each bin. **D** Half-lives for genes binned by median poly(A) tail length. Numbers indicate the number of genes in each bin. **E** 3’UTR length for genes binned by median poly(A) tail length. Numbers indicate the number of genes in each bin. **F** 3’UTR length of proximal and distal 3’UTR isoforms for genes with 2 annotated 3’UTR isoforms binned by median poly(A) tail length for each isoform. Numbers indicate the number of 3’UTR isoforms in each bin. **G** Median poly(A) tail length per gene for lncRNA genes in HeLa S3 fractionation experiments (**left**) and mouse brain fractionations (**right**). **H** Median poly(A) tail length per gene between cytoplasmic and nuclear fractions for mouse brain fractionation experiments. **I** Median poly(A) tail length per gene compared across fractions from HeLa S3 cell lines for ribosomal protein genes (‘Ribosomal’), immediate early genes (‘IEGs’) and lncRNAs. **J** Cytoplasmic-to-nuclear enrichment of genes quantified by ENCODE for HeLa cell lines versus FLAM-seq quantification.

**Supplementary Fig. 6.**
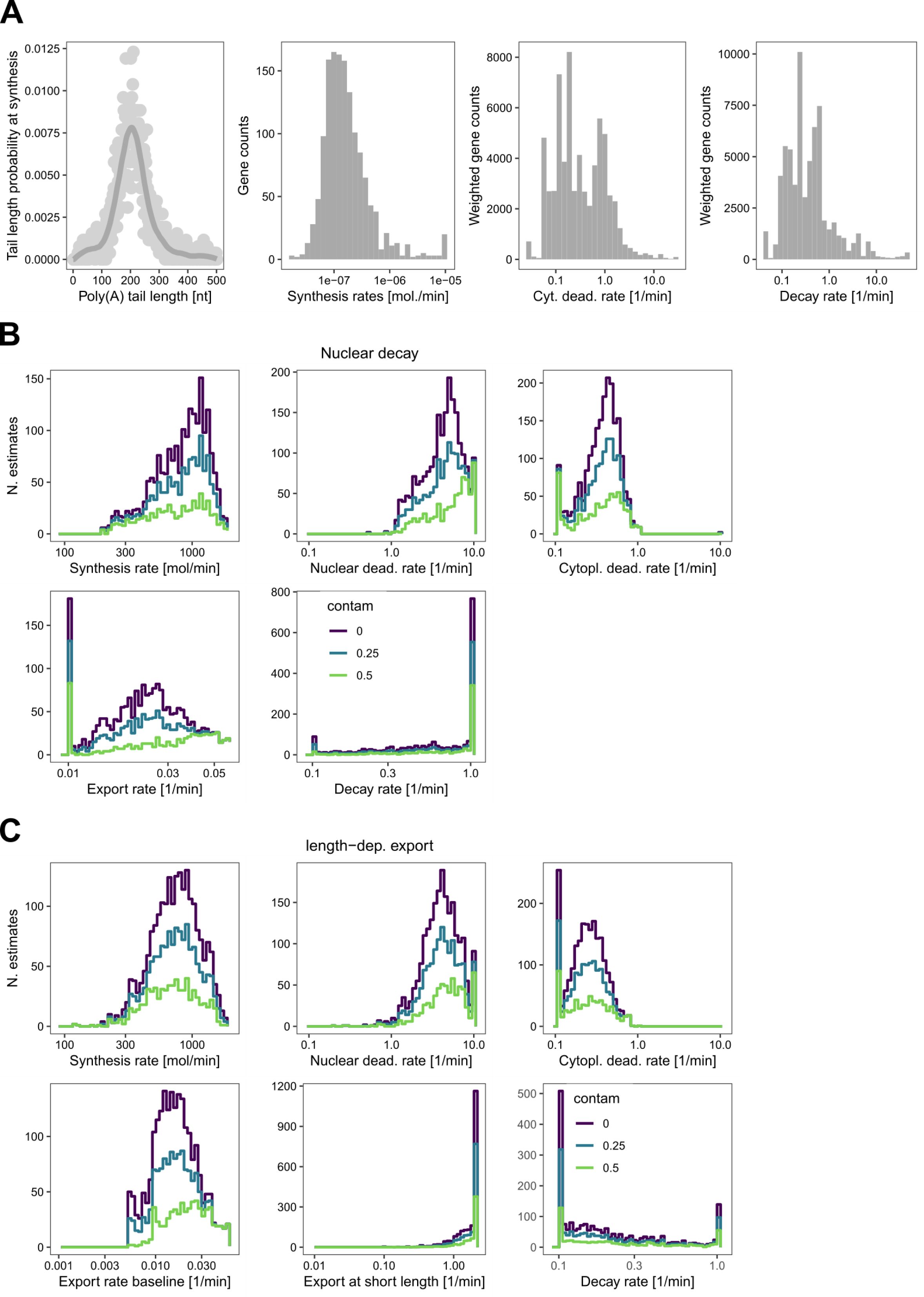
A mathematical model predicts fast nuclear deadenylation. **A** Data used for initial parameter guesses in the optimization: slp parameter (left) is estimated from intronic reads of 12 FLAM-seq fractionation replicates; synthesis rates are estimated by summing up all transcription rates estimated from Eisen et al. (2020) (distribution per gene shown in middle left), nuclear and cytoplasmic deadenylation rates are initialized according to the mean of the deadenylation rates measured by Eisen et al. (2020) (distribution weighted for gene expression in FLAM-seq data shown in middle right); decay rate: same as before (right). **B** From top left to bottom right: final distributions from optimization of synthesis rate, nuclear and cytoplasmic deadenylation rate, export rate, decay rate. Parameters were estimated with cytoplasmic contamination of nuclear RNA set at 0, 25 and 50% (see color legend), for the nuclear decay - constant export (ndce) model. **C** Same as B., for the no nuclear decay, tail length dependent export (nndle) model.

**Supplementary Fig. 7.**
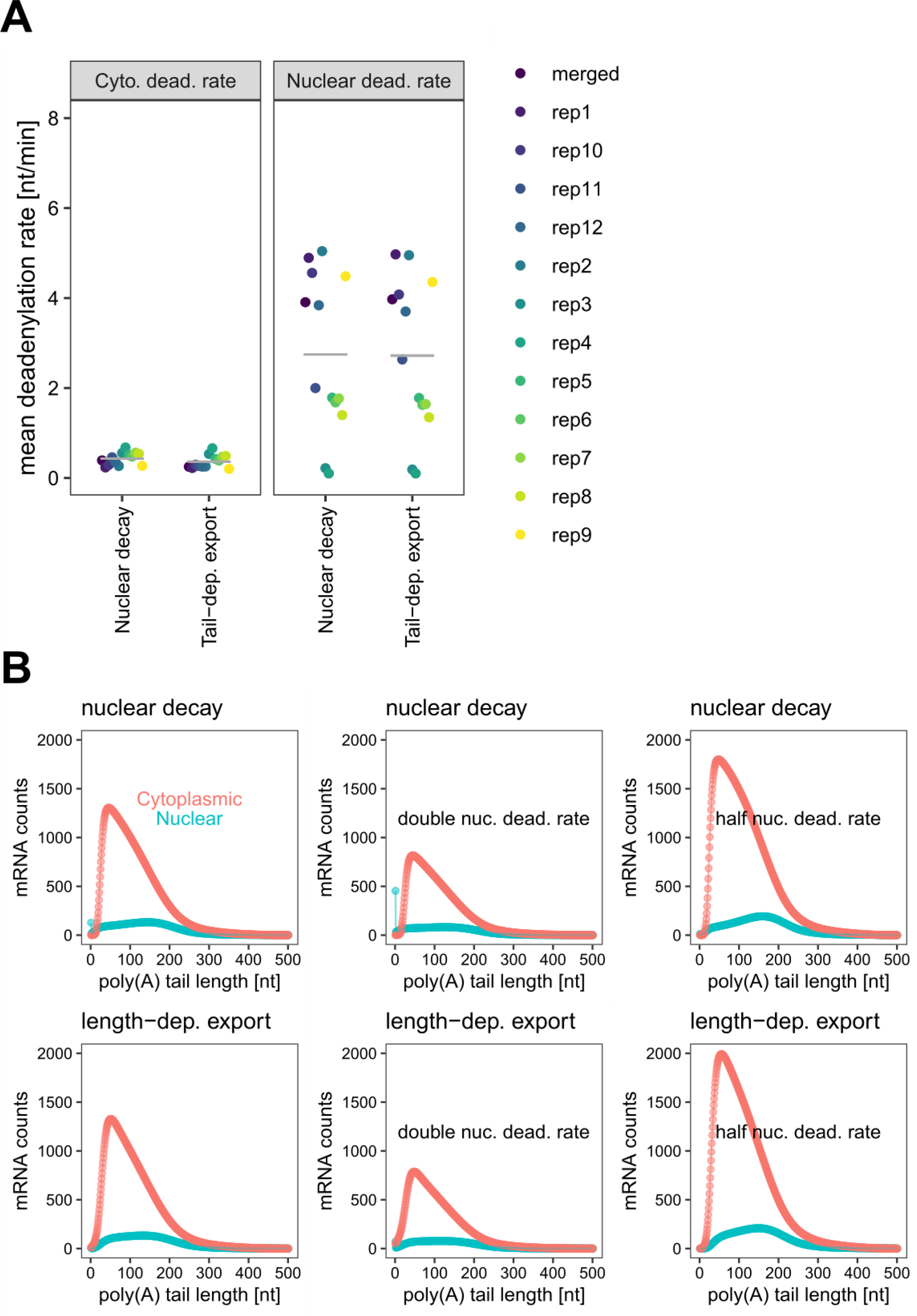
A mathematical model predicts fast nuclear deadenylation. **A** Values of k_1_ and k_2_ (nuclear and cytoplasmic deadenylation rate) estimated independently from 12 fractionation experiments for both nuclear decay constant export (ndce) and no nuclear decay, tail length dependent export (nndle). k_1_ values are consistently higher than k2, except for two replicates, which were performed in the same batch, and where tail lengths were overall longer than for all other replicates (possibly a technical artifact of the poly(A) selection step in the FLAM-seq protocol as discussed in the results section). **B** Simulation of mRNA levels in the nucleus (green) and cytosol (red) with the model assuming constant export and nuclear decay (ndce model; top) and the model assuming tail-length dependent export (nndle model; bottom), using the average of the estimated parameters distributions as input (left), or increasing and decreasing the nuclear deadenylation k_1_ rate 2-fold (middle and right). For this specific example, we assumed a cytoplasmic contamination in the nuclear fraction of 0%.

**Supplementary Table 1.** Definition and description for all parameters used in the ndce and nndle models optimization. Means of initial guesses are shown, while esimated values are shown as mean and standard deviation of merged distribution from 1,000 iterations for 0, 25 and 50% cytoplasmic contamination of nuclear fraction, for ndce and nndle (comma separated).

